# Intracranial Human Recordings Reveal Intensity Coding for the Pain of Others in the Insula

**DOI:** 10.1101/2021.06.23.449371

**Authors:** Efe Soyman, Rune Bruls, Kalliopi Ioumpa, Laura Müller-Pinzler, Selene Gallo, Elisabeth C.W. van Straaten, Matthew W. Self, Judith C. Peters, Jessy K. Possel, Yoshiyuki Onuki, Johannes C. Baayen, Sander Idema, Christian Keysers, Valeria Gazzola

## Abstract

Based on neuroimaging data, the insula is considered important for people to empathize with the pain of others, whether that pain is perceived through facial expressions or the sight of limbs in painful situations. Here we present the first report of intracranial electroencephalographic (iEEG) recordings from the insulae collected while 7 presurgical epilepsy patients rated the intensity of a woman’s painful experiences viewed in movies. In two separate conditions, pain was deduced from seeing facial expressions or a hand being slapped by a belt. We found that broadband activity in the 20-190 Hz range correlated with the trial-by-trial perceived intensity in the insula for both types of stimuli. Using microwires at the tip of a selection of the electrodes, we additionally isolated 8 insular neurons with spiking that correlated with perceived intensity. Within the insula, we found a patchwork of locations with differing selectivities within our stimulus set, some representing intensity only for facial expressions, others only for the hand being hit, and others for both. That we found some locations with intensity coding only for faces, and others only for hand across simultaneously recorded locations suggests that insular activity while witnessing the pain of others cannot be entirely reduced to a univariate salience representation. Psychophysics and the temporal properties of our signals indicate that the timing of responses encoding intensity for the sight of the hand being hit are best explained by kinematic information; the timing of those encoding intensity for facial expressions are best explained by shape information in the face. In particular, the furrowing of the eyebrows and the narrowing of the eyes of the protagonist in the movies suffice to predict both the rating of and the timing of the neuronal response to the facial expressions. Comparing the broadband activity in the iEEG signal with spiking activity and an fMRI experiment with similar stimuli revealed a consistent spatial organization for the representation of intensity from our hand stimuli, with stronger intensity representation more anteriorly and around neurons with intensity coding. In contrast, for the facial expressions, we found that the activity at the three levels of measurement do not coincide, suggesting a more disorganized representation. Together, our intracranial recordings indicate that the insula encodes, in a partially intermixed layout, both static and dynamic cues from different body parts that reflect the intensity of pain experienced by others.

## 1. Introduction

Sharing the distress of others is central to empathy. fMRI studies show that a number of brain regions involved in the direct experience of pain also increase their activity while participants perceive the pain of others, including the cingulate cortex, the insula, and the somatosensory cortices (Jauniaux et al., 2019; Keysers et al., 2010; Lamm et al., 2011; Timmers et al., 2018). Across humans, primates, and rodents, lesions in these regions impair the perception and the sharing of others’ emotions (Paradiso et al., 2021). Directly recording electrical signals from these regions in humans would complement the more indirect fMRI measurements and sharpen our understanding of how these regions represent the intensity of other people’s pain, for at least two reasons. First, fMRI records a mixed signal that includes synaptic input and local neural processing. Localizing BOLD activity that encodes a particular task or stimulus property in a particular brain region thus cannot ensure that neurons in that region actually have spiking activity that encodes that property (Boynton, 2011). For instance BOLD signals in V1 fluctuate based on whether a stimulus is perceived or not in binocular rivalry (Boynton, 2011; Maier et al., 2008). In contrast, simultaneous electrical recordings show that broadband gamma activity, which is tightly coupled to spiking (Bartoli et al., 2019; Buzsáki et al., 2012; Miller et al., 2014), in the same region responds to a stimulus equally well whether it is perceived or suppressed - only the slower components, <20Hz, that are known to carry feedback synaptic input, fluctuate with perception (Maier et al., 2008). Being able to record electrical activity, particularly in the broadband gamma range, is thus critical to localize where in this circuitry neuronal spiking indeed represents the pain of others. Second, fMRI’s low temporal resolution makes it difficult to characterize the time-course of responses.

For the anterior cingulate we have intracranial recordings: Hutchison (1999) documented a single neuron in epileptic patients that responded to the sight of a finger being pin-pricked with increased firing rate, and a recent rodent study revealed that cingulate neurons responding to pain experience have responses that increase with the intensity of the pain experienced by another rat (Carrillo et al., 2019). In contrast, although the insula is central in the neuroimaging literature on empathy, and shows increases of BOLD signal for watching painful compared to non-painful social stimuli (Jabbi et al., 2007; Jauniaux et al., 2019; Lamm et al., 2011; Meffert et al., 2013; Singer et al., 2004; Timmers et al., 2018; Wicker et al., 2003), we still lack such intracranial recordings while individuals witness the pain of others. Intracerebral EEG (iEEG) has been recorded in the insula during the self-experience of pain (e.g., Liberati et al., 2020), and the insula and adjacent SII are the only cortical region where iEEG electrode stimulation can induce painful sensations (Jobst et al., 2019; Mazzola et al., 2012), but to our knowledge there are no published studies recording from insular electrodes while patients witness the pain of others. The degree to which neuronal activity local to the insula, as opposed to feedback synaptic input from other regions such as the cingulate, encodes the intensity of other people’s pain therefore remains unclear and the time-course of such neural activity remains under characterized.

To fill this gap and characterize the electrophysiological responses of the insula to the pain of others, we collected depth electrode recordings from 7 epileptic patients during pre-surgical exploration, while they rated the different intensities of pain they perceived another person in a video to experience (Fig. 1a,b). All these patients had macro electrodes in their insulae that yielded local field potentials (LFP) capable of measuring broadband gamma activity (circles in Fig. 1c). Three patients, additionally, had micro electrodes at the tip of some macro electrodes to record from isolated insular neurons (pluses in Fig. 1c). Our stimuli included two ways in which pain is perceived in others (Fig. 1a). Half the stimuli (Faces) showed a female receiving electroshocks on the hand and expressing pain through facial expressions (furrowing eyebrows and tightening eyes). The other half (Hands) showed the protagonist’s hand slapped by a leather belt, and pain intensity had to be deduced from the movements of the belt and the hand. In both cases, movies, rather than static images, were chosen to provide richer and more ecologic stimuli and provide information about the temporal dynamics with which such movies are represented in a field still dominated by the presentation of static images (Adolphs et al., 2003; Zinchenko et al., 2018). We used these two classes of stimuli, because both tap into the visual perception of other people’s pain, and we start to understand that they do so through dissociable routes (Jauniaux et al., 2019; Keysers et al., 2010; Timmers et al., 2018). For instance, the hand stimuli depend on the hand region of the somatosensory cortex (Gallo et al., 2018), while facial expressions are depend on the ventral somatosensory cortex and the insula (Adolphs et al., 2000; Dal Monte et al., 2013; Mattavelli et al., 2019). Based on fMRI studies showing increased activity in the insula for more painful stimuli, we hypothesized increases in power in higher LFP frequencies and spike counts for higher pain intensity ratings, thus used one-tailed testing unless specified otherwise.

**Figure 1.**
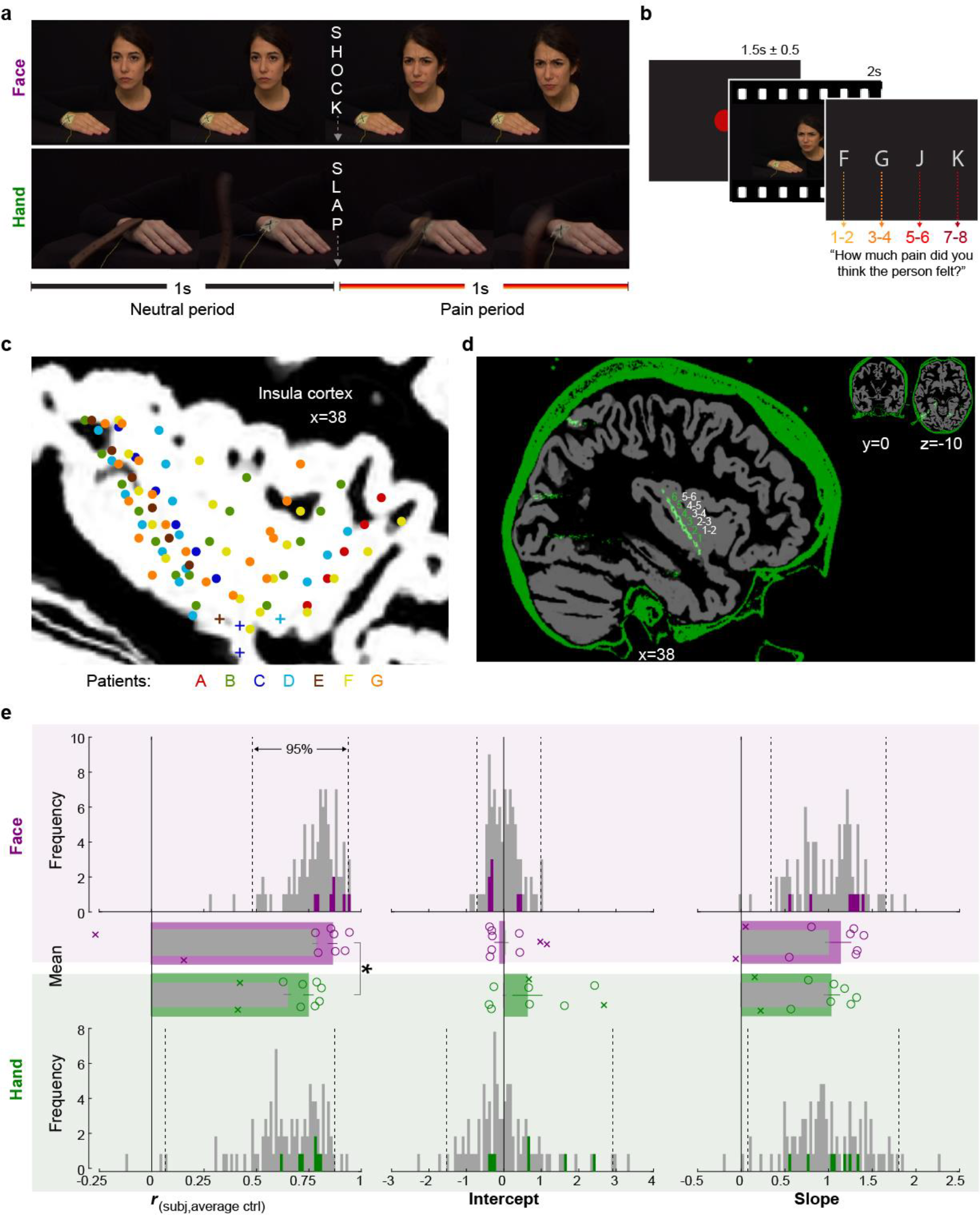
Experimental design, recording site locations, and behavioral pain ratings. **(a)** Frames extracted from a Hand and a Face movie. For the Face, the first second of each movie showed a neutral facial expression, the second, the facial reaction to the shock. For the Hand, the movie started with the belt resting on the hand. The first second showed the belt lifting and coming down again, to hit the hand at the 1 s mark exactly. The hand then reacted to the force of the belt. Both the slap and the shock delivery happened in the middle of the movies, splitting them into a 1 s neutral and a 1 s pain period. **(b)** Single trial structure diagram. After the presentation of each video, patients expressed their choice at their pace using the keyboard keys f, g, j, k for pain intensities 1-2, 3-4, 5-6, 7-8, respectively. ITI started with participant’s response. **(c)** Position (i.e., the midpoint between two adjacent electrodes) of the 85 bipolar macro electrode recording sites shown as dots and of the micro electrode locations shown as plusses, color coded by patient. Data from the two hemispheres and all latero-medial coordinates are projected here onto a single sagittal slice of the insula taken at X=38 from the brain of one of the patients. For a list of all MNI coordinates, see Supplementary File 1. **(d)** Graphical illustration of how a bipolar recording for one patient and one insular electrode was computed. In green the CT and in gray the T1 scan from Patient C. The annular structures along the electrode shaft in the CT correspond to individual macro electrode contacts (green 1, 2, 3…). Recordings from adjacent pairs of contacts along the electrode were subtracted to calculate bipolar recordings (white 1-2, 2-3…). **(e)** From left to right, Spearman correlation coefficient r, intercept, and slope values from the linear regression for Hand (green) and Face (purple). Histograms: values for the control group illustrate the similarity between the ratings of each participant in the control group and the average of the other controls, and are shown as gray; the similarity between each of 7 patients with the average of the control group are shown in colors. Dotted lines mark the 2.5% and 97.5% of the control group. Bar graphs: Mean±SEM of the controls (gray) and the 7 included patients (color) with individual patients as circles. In the bar graphs, we also show as Xs the corresponding behavioral performance metrics of the two patients that were excluded due to atypical use of the response keys. These patients were not included in the Mean and SEM calculations.

## 2. Results

### 2.1. The Pain Intensity Ratings of Patients were within the Normal Range

To assess whether the behavior of the patients was representative of the general population, we compared patients’ (3 males, 4 females, 34.3y±9std) ratings with those of 93 healthy volunteers (54 females, 32.7y±9std, Table 2), who took part in an online version of the video pain rating task. Table 1 shows the distribution of ratings separately for patients and on average for controls. We calculated three metrics of similarity between the ratings of the patients and the control group: the Spearman’s rank order correlation, the slope, and the intercept of a simple linear regression between each patient’s ratings and the average ratings of the control sample (Fig. 1e). The patients revealed correlation coefficients, slopes, and intercepts (green and purple bars) within the 2.5 and 97.5 percentiles of the corresponding control sample distributions (gray bars), except for one correlation coefficient for Faces, where a patient rated the Faces with unusually high concordance with the average of the control groups. This verified that these 7 patients were not impaired in their ability to rate intensity from our videos. In both the patient and the control sample, we observed that correlation coefficients for Faces were significantly greater than for Hands (Patient: *t*_*(6)*_=3.81, *p*_*2*_=0.009, BF_10_=7.39; Control: *W*=3895, *p*_*2*_=10^−12^, BF_10_=3×10^5^, Fig. 1e), suggesting more interpersonal agreement in pain intensity ratings for Faces than Hands. In contrast to the higher agreement for Faces, the average rating was slightly higher for Hands than Faces in both the patient and the control sample (Patient: *t*_*(6)*_=2.6, *p*_*2*_=0.04, BF_10_=2.3; Control: *W*=1116, *p*_*2*_=0.0005, BF_10_=143). Taken together, these findings indicate that the intensity rating behavior of the patient samples were similar to the patterns observed in the healthy population.

**Table 1.** Pain ratings in patients and controls. **Left:** Number of trials (out of the 60 Hand and 60 Face trials for Patients, or 30 and 30 for Controls) per rating per participant for the Hand and Face conditions. For the age and gender matched control group, only the average across the 93 controls is shown. **Middle**: Mean (M) rating for the Hand or Face. Our patients reported slightly higher pain intensity ratings for our Hand than Face stimuli (*t*_*(6)*_=2.6, *p*_*2*_=0.04, BF_10_=2.3), the same was true for the age- and gender-matched controls (n=93, *W*=1116, *p*_*2*_=0.0005, BF_10_=143). This was somewhat surprising, because the Hand and Face stimuli were rated as similarly intense in a validation study that preceded stimulus selection (Gallo et al., 2018). Age- and gender-matched controls also rated the Hand stimuli as slightly more intense, although the difference was less pronounced. **Right:** Standard deviation of the ratings for each participant. Because the efficiency of a regression depends on the standard deviation of the predictor, and much of our results depend on the relation between rating and iEEG responses, we calculated the standard deviation for each participant and condition. The standard deviations were normally distributed (all Shapiro-Wilk *p*>0.25), we then used a t-test to compare them across the two conditions. We found no significant difference amongst the patients (*t*_*(6)*_=-1.442, *p*_*2*_=0.199, BF_10_=0.747). Differences we find in the correlations between rating and iEEG across Hand and Face stimuli therefore cannot be due to difference in the efficiency of these two estimations.

|  |  | Rating |  |  |  |  |  |  |  | M (Rating) |  | SD (Rating) |  |
| --- | --- | --- | --- | --- | --- | --- | --- | --- | --- | --- | --- | --- | --- |
|  |  | Hand |  |  |  | Face |  |  |  | Hand | Face | Hand | Face |
|  |  | 1-2 | 3-4 | 5-6 | 7-8 | 1-2 | 3-4 | 5-6 | 7-8 |  |  |  |  |
| Patient | A | 1 | 23 | 26 | 10 | 24 | 17 | 15 | 4 | 2.75 | 1.98 | 0.75 | 0.97 |
|  | B | 0 | 10 | 30 | 20 | 22 | 21 | 8 | 9 | 3.17 | 2.07 | 0.69 | 1.06 |
|  | C | 0 | 0 | 26 | 34 | 34 | 26 | 0 | 0 | 3.57 | 1.43 | 0.50 | 0.50 |
|  | D | 1 | 23 | 23 | 13 | 21 | 25 | 14 | 0 | 2.80 | 1.88 | 0.80 | 0.76 |
|  | E | 17 | 17 | 19 | 7 | 21 | 12 | 17 | 10 | 2.27 | 2.27 | 1.01 | 1.12 |
|  | F | 15 | 24 | 15 | 6 | 23 | 19 | 15 | 3 | 2.20 | 1.97 | 0.94 | 0.92 |
|  | G | 23 | 14 | 16 | 7 | 22 | 17 | 15 | 6 | 2.12 | 2.08 | 1.06 | 1.01 |
|  | Mean | 8 | 16 | 22 | 14 | 24 | 20 | 12 | 5 | 2.70 | 1.95 | 0.82 | 0.91 |
| Control | Mean | 10 | 12 | 6 | 2 | 13 | 10 | 5 | 2 | 2.02 | 1.84 | 0.74 | 0.81 |

**Table 2:**
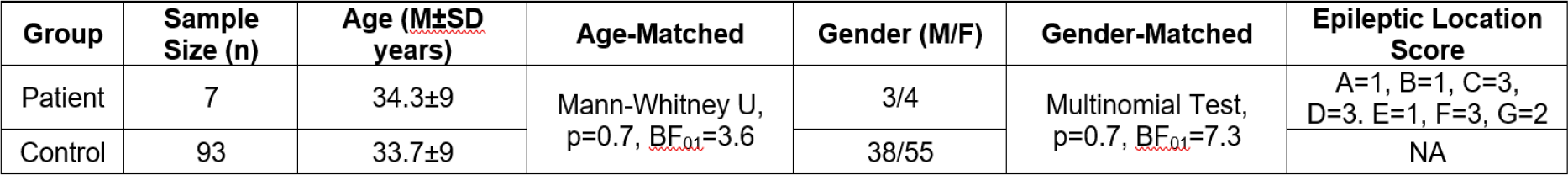
Participants’ demographics and epileptic status. Our 7 patients were matched in Age and Gender to the online control group from which we obtained normative movie ratings. The last column indicates for our patients, their post-operative status. Three patients had other brain regions than insula surgically removed, and afterwards had no more attacks (marked with 1), suggesting that the foci were clearly outside the insula. One patient had a region other than the insula surgically removed, because the monitoring had suggested that the foci was outside of the insula, however, the patient continued to have post-surgical attacks (marked with 2). For three patients, no surgery was performed because there was no clear link between electrode locations and epileptic attacks (marked with 3).

| Group | Sample Size (n) | Age (M±SD years) | Age-Matched | Gender (M/F) | Gender-Matched | Epileptic Location Score |
| --- | --- | --- | --- | --- | --- | --- |
| Patient | 7 | 34.3±9 | Mann-Whitney U, p=0.7, $BF_{01}$=3.6 | 3/4 | Multinomial Test, p=0.7, $BF_{01}$=7.3 | A=1, B=1, C=3, D=3, E=1, F=3, G=2 |
| Control | 93 | 33.7±9 | | 38/55 | | NA |

### 2.2. LFP Activity in the Insula Correlates with the Perceived Intensity of the Pain of Others

Plotting the power over the bipolar electrodes as a function of perceived intensity, irrespectively of whether Hand or Face videos were shown, suggests an increase in power for the highest ratings (Fig. 2d). Correlating power with reported intensity and applying cluster corrections revealed a cluster of positive correlations ranging from 20-190 Hz and 1.12-1.62 s (*p*_*1*_<0.001; *p*_*1*_=one-tailed p value), another cluster of positive correlations at very low frequencies (1-6 Hz, 0.02-2.055 s; *p*_*1*_<0.001), and a small cluster of negative correlations (13-17 Hz, 1.295-1.83 s; *p*_*1*_=0.004, not further discussed; Fig. 2a). Intensity coding was apparent in all traditional frequency ranges, except alpha (Fig. 2b), and, as expected, was significant in the pain period. With no obvious differences among frequency-bands above alpha, we henceforth used the frequency band 20-190 Hz for all analyses and refer to it as broadband power (BBP). We concentrate on BBP rather than oscillatory signals in lower frequencies because BBP is more closely linked to neural spiking (Bartoli et al., 2019; Buzsáki et al., 2012; Miller et al., 2014), cannot be explored in non-invasive EEG recordings, and is the frequency range that can supplement the information available for the substantial fMRI literature (Boynton, 2011; Maier et al., 2008). The temporal profile of the BBP-rating association revealed two periods with significant positive correlations: 1.1375–1.54 s and 1.7375–1.9575 s (Fig. 2b). Averaging BBP power over the entire pain period revealed that out of 85 macro contacts, 27 (32%) showed a significant positive correlation (assessed as *p*_*1*_<0.05, Fig. 2c) between perceived intensity and BBP (n=120 trials, all *r*_*S(118)*_>0.156, p_1_<0.045), which was extremely unlikely to occur by chance (Binomial *p*_*1*_=5×10^−15^, BF_+0_=3×10^12^). Furthermore, randomly picking 85 electrodes anywhere in the brain yielded BBP-rating associations that were significantly lower than those we found in the insula (*p*_*1*_=4×10^−5^, Fig. 2g), confirming that the BBP in the insula has enriched intensity coding. Splitting trials based on reported intensity and identifying moments in which the intensity coding is significant in an ANOVA confirmed that BBP scaled with pain ratings from 1.0925 to 1.6975 s (Fig. 2e). Averaging the BBP over the 1 s neutral and 1 s pain period and using a period (neutral, pain) x rating repeated-measures ANOVA revealed a significant interaction effect (*F*_*(2*.*445,205*.*348)*_=37.49, *p*=8×10^−17^, BF_incl_=85925, Fig. 2f). Planned comparisons show BBP was similar during the neutral and the pain periods for videos rated 1-2 (*W*=1903, *p*_*2*_=0.742, BF_10_=0.126) and 3-4 (*W*=1801, *p*_*2*_=0.909, BF_10_=0.146), but was higher during the pain than neutral period when rated 5-6 (*t*_*(84)*_=3.42, *p*_*2*_=0.001, BF_10_=24.31) and 7-8 (*t*_*(84)*_=7.29, *p*_*2*_=2×10^−10^, BF_10_=6×10^7^). BBP was similar during pain periods rated 1-2 and 3-4 (*W*=1966, *p*_*2*_=0.545, BF_10_=0.120), while BBP increased in the pain period for further rating increases (3-4 vs 5-6: *W*=1065, *p*_*2*_=8×10^−4^, BF_10_=15.206; 5-6 vs 7-8: *W*=309, *p*_*2*_=3×10^−11^, BF_10_=950944). These results indicate that the effect of reported intensity depends mainly on BBP power increases for the two highest intensity ratings.

**Figure 2.**
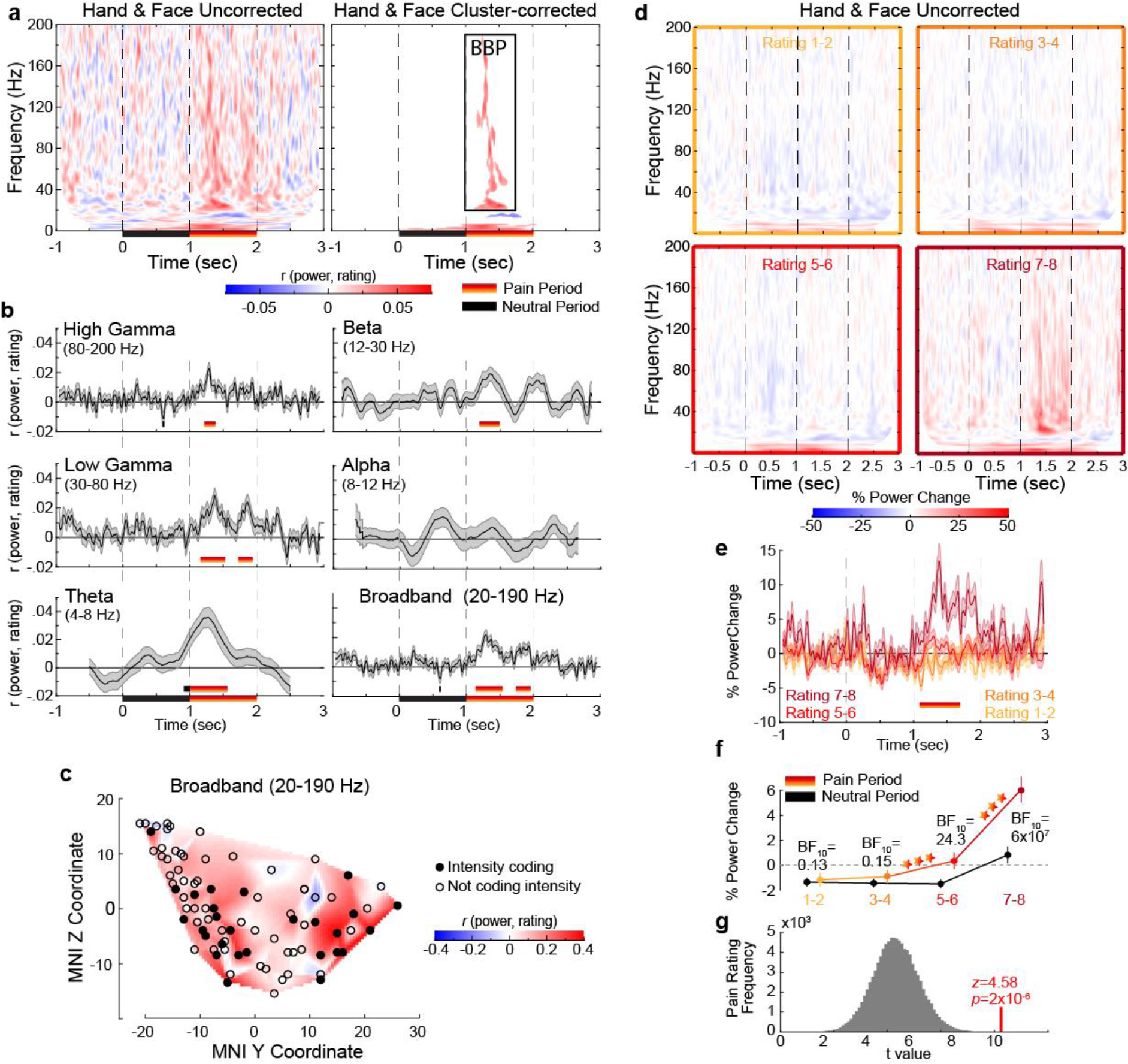
Intensity coding in the insula LFP activity for Hands and Faces together. **(a)** For each frequency and time relative to stimulus onset, the average *r*_*S*_ value over all insular bipolar recordings between iEEG power and rating for Face and Hand trials together, without (left) and with cluster correction for multiple comparisons (right). BBP: Broadband power, the cluster of significant positive intensity coding frequencies (i.e., *r*_*S*_>0, 20-190 Hz) used throughout the paper. **(b)** Mean±SEM time course of intensity coding in different frequencies and BBP over the 85 recordings when Face and Hand trials are combined. Above the x-axis, black and yellow-to-red bars show periods of significant intensity coding after circular shift correction for multiple comparisons during the neutral and pain periods, respectively. Below the x-axis, the black bar marks the neutral and the yellow-to-red bar indicates the pain period. **(c)** Intensity coding in the 85 bipolar recordings is shown as significant (*p*_*1*_<0.05, filled black circles) or non-significant (*p*_*1*_>0.05, open circles) based on the MNI y (anterior-posterior) and z (dorso-ventral) coordinates. The heatmap shows the interpolated intensity coding value between these locations. Electrodes in the right and left insula are projected onto the same sagittal representation. **(d)** Mean percent power changes relative to the baseline period (1 s before the onset of videos) across all 85 bipolar recordings as a function of time and frequency for all trials rated 1-2 (top left), 3-4 (top right), 5-6 (bottom left), and 7-8 (bottom right) when Hand and Face trials analyzed together. Note the pronounced increase of power in a wide frequency range spanning from 20-190 Hz. **(e)** Mean±SEM percent power change time courses averaged in the broadband range (20-190 Hz), separately for each reported rating. **(f)** Mean±SEM percent power change values as a function of reported intensity separately for the neutral (black) and pain (yellow-to-red) periods when combining Hand and Face trials. BF_10_ values: Bayes-factor quantifying evidence for H_1_ relative to H_0_ from a non-parametric t-test comparing BBP power during the pain period against that during the neutral period. ***: *p*<0.001 relative to the preceding reported intensity. **(g)** The t value of a t-test comparing the intensity coding of all 85 bipolar recordings combining Hand and Face trials within the pain period (1–2 s post-stimulus onset) in the insula against zero (red) was higher than the distribution of the corresponding t values obtained when performing the same test using 85 bipolar recordings randomly selected 100,000 times from the macro electrode contacts of our 7 patients anywhere in the brain.

### 2.3. Intensity Coding Arises Earlier for Hands than Faces

To investigate how intensity coding depends on the stimulus, we then separated Face and Hand trials (Fig. 3e). For Hands, there was a cluster of positive power-rating correlations between 33 and 145 Hz and 1.1375 and 1.53 s (*p*_1_<0.001), one at very low frequencies (1–6 Hz, 0.3475– 1.9275 s; *p*_1_=0.002), and a small negative correlation cluster (1–4 Hz, -0.5–0.175 s; *p*_1_=0.002) that survived the cluster correction (Fig. 3a,b). For Faces, there was a trend for a large cluster of positive correlations between 1 and 7 Hz and 0.975 and 2.3475 s (*p*_1_=0.028 at alpha=0.025, Fig. 3a,b). A direct comparison of the time-frequency plots revealed regions of intensity coding earlier for Hands than for Faces (Fig. 3c,d). The analysis of the time courses of the frequency bands that showed the maximal positive correlation clusters for Hands and Faces separately indicated a significant positive correlation period during the pain period for Hands, but no effect for Faces (Fig. 3f). The absence of significant intensity coding for Faces in the time-frequency decomposition and the time course analysis was due to a lack of power after correction for multiple comparisons, and should therefore be seen as purely explorative. Focusing on the more powerful BBP range of interest (20-190 Hz) identified independently of stimulus type (Fig. 2a), we found significant intensity coding for the Hand from 1.0075 to 1.4375 s (hereafter called Early Period) and for the Face from 1.75 to 1.8625 s and from 1.905 to 1.975 s (jointly called Late Period, Fig. 3g,h). The insula thus reflects in broadband activity the perceived intensity with differential time courses for the Hand and Face videos in the current study. To explore the shape of the BBP-rating relation, we averaged BBP over time for the early and the late periods for each pain rating separately (Fig. 3h). For the early period, a stimulus (Hand, Face) x rating repeated-measures ANOVA revealed a significant interaction (Greenhouse-Geisser-corrected *F*_(2.183,102.621)_=13.55, *p*=3×10^−6^, BF_incl_=2×10^6^). Planned comparisons provided evidence that BBP for Faces in the Early Period was similar for consecutively increasing painfulness level pairs (n=48 since all possible rating options were used by 4 patients with a total of 48 electrodes, all *t*_(47)_<0.252, *p*_*2*_>0.802, BF_10_<0.163), whereas, there was an orderly increase in broadband power for increasing pain ratings for Hands from 3-4 onwards (3-4 vs 5-6: *t*_(47)_= 5.97, *p*_*2*_=3×10^−7^, BF_10_= 51110; 5-6 vs 7-8: *W*= 188, *p*_*2*_=2×10^−5^, BF_10_= 764.63). However, BBP for ratings of 1-2 was unexpectedly higher compared to the ratings of 3-4 (*W*=1014, *p*_*2*_=3×10^−6^, BF_10_= 847.14). A similar ANOVA for the Late Period, revealed evidence for the absence of an interaction (*F*_(3,141)_=0.55, *p*=0.650, BF_incl_=0.034). There was only a significant main effect of rating (*F*_(3,141)_=16.54, *p*=3×10^−9^, BF_incl_=2×10^7^), indicating that BBP in the Late Period of the Hand and Face videos together was the same for ratings 1-2 and 3-4 (*W*= 597, *p*_*2*_=0.931, BF_10_= 0.163), but thereafter showed significant increases with each consecutive increase in pain ratings (3-4 vs 5-6: *t*_(47)_= 3.46, *p*_*2*_=0.001, BF_10_= 25.147; 5-6 vs 7-8: *t*_(47)_= 2.90, *p*_*2*_= 0.006, BF_10_= 6.292). Taken together, these analyses indicate BBP in the insula reflects perceived intensity only for the Hand stimuli in the Early, and for both stimulus types in the Late Period.

**Figure 3.**
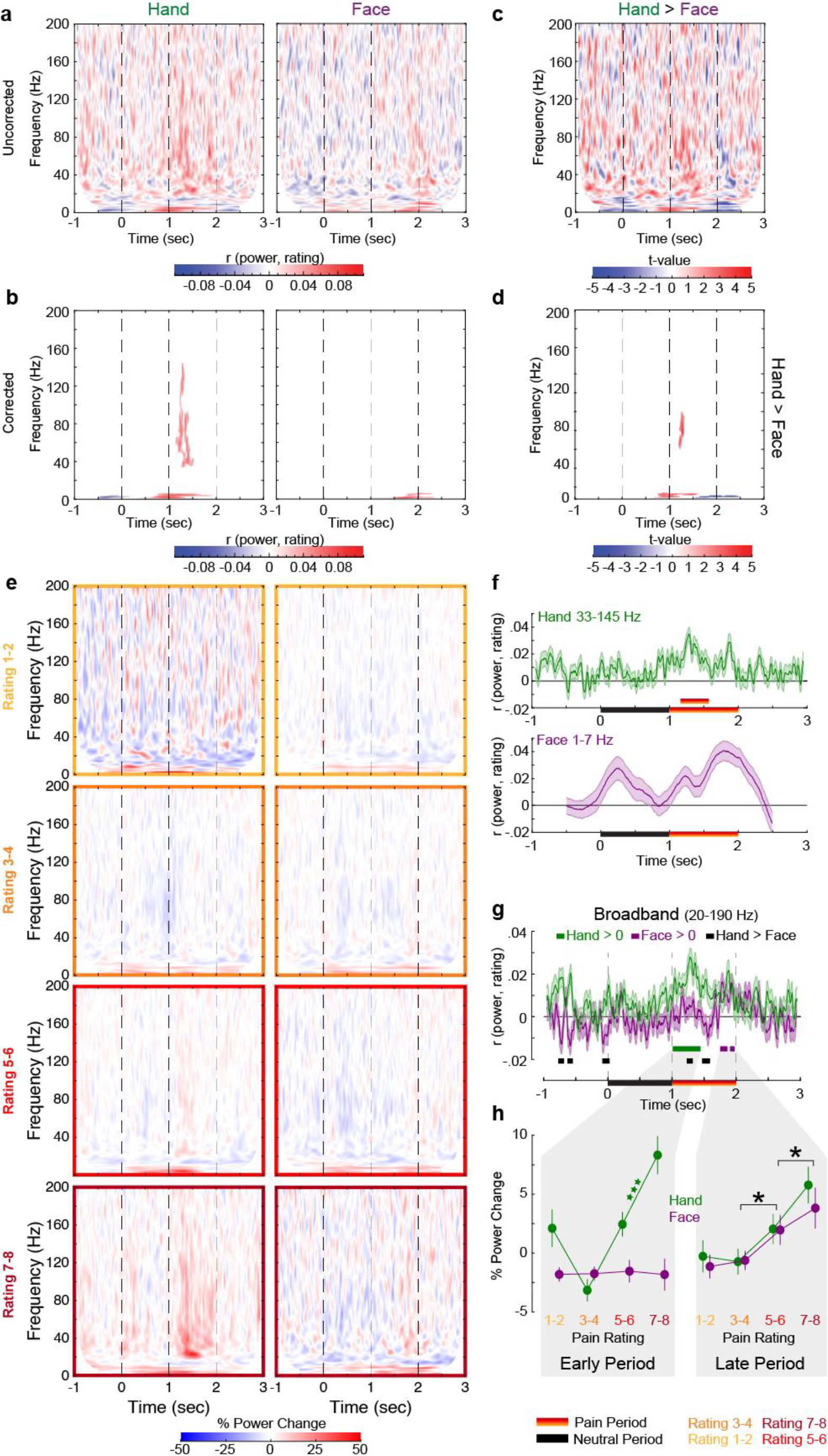
Intensity coding in the insula LFP activity for Hands and Faces separately. **(a)** For each frequency and time relative to stimulus onset, the average *r*_*S*_ value over all insular bipolar recordings between iEEG power and rating for Hand (left) and Face (right) trials separately. **(b)** As in (a), but cluster corrected for multiple comparisons. **(c)** For each frequency and time relative to stimulus onset, t values of paired-samples t-tests comparing the Spearman correlations obtained for the Hands and Faces. **(d)** As in (c), but cluster corrected for multiple comparisons **(e)** Mean percent power changes relative to the baseline period (1 s before the onset of videos) across all 85 bipolar recordings as a function of time and frequency for all trials rated 1-2 (first row), 3-4 (second row), 5-6 (third row) and 7-8 (fourth row) for the Hand (left) and Face (right) trials separately. **(f)** Mean±SEM time courses of intensity coding for the largest positive correlation cluster (frequency range as indicated) for Hands (top, 33-145 Hz) and Faces (bottom, 1-7 Hz) separately. The yellow-to-red bar indicates a correlation coefficient period significantly greater than 0 after circular shift cluster correction for multiple comparisons. **(g)** Mean±SEM time course of intensity coding in BBP (20-190 Hz) for Hands and Faces separately. *r*_*S*_>0 indicated with green bars for Hands and purple bars for Faces. Black bars indicate *r*_*S_*Hand_>*r*_*S_Face*_. The early and late periods that result for Hands and Faces, respectively, are used throughout the paper. **(g)** Mean±SEM percent power change in the broadband frequency rands as a function of rating for Hands and Faces in the early and late periods separately. Green ***: *p*<0.001 relative to the preceding intensity for the Hand. Black *: *p*<0.001 main effect of rating (i.e., combining Hand and Face). Purple BF_10_: evidence for a lack of difference across ratings for the Hand. Black BF_10_: lack of difference between rating 1-2 and 3-4 when Hand and Face trials combined.

### 2.4. The Timing of Shape Information Matches that of Face Intensity Coding

Having observed critical differences in the temporal profiles of intensity coding for Hands and Faces, we next assessed whether these differences could arise from the timing of different pain intensity cues depicted in the two video types. We first subjected our stimuli to more detailed, time-resolved analyses to describe the temporal evolution of the the motion information in the Hand videos, and the motion and the shape information in the Face videos. Motion information for both Hands and Faces was quantified based on pixel-based intensity changes across consecutive frame pairs. Shape information for Faces was estimated using an automated face analysis software to extract the two most reliable shape features of painful facial expressions, how lowered the eye-brows and how tightened the eye-lids are (facial Action Units AU4 and AU7, respectively, Fig. 4a, Kunz et al., 2019)

**Figure 4.**
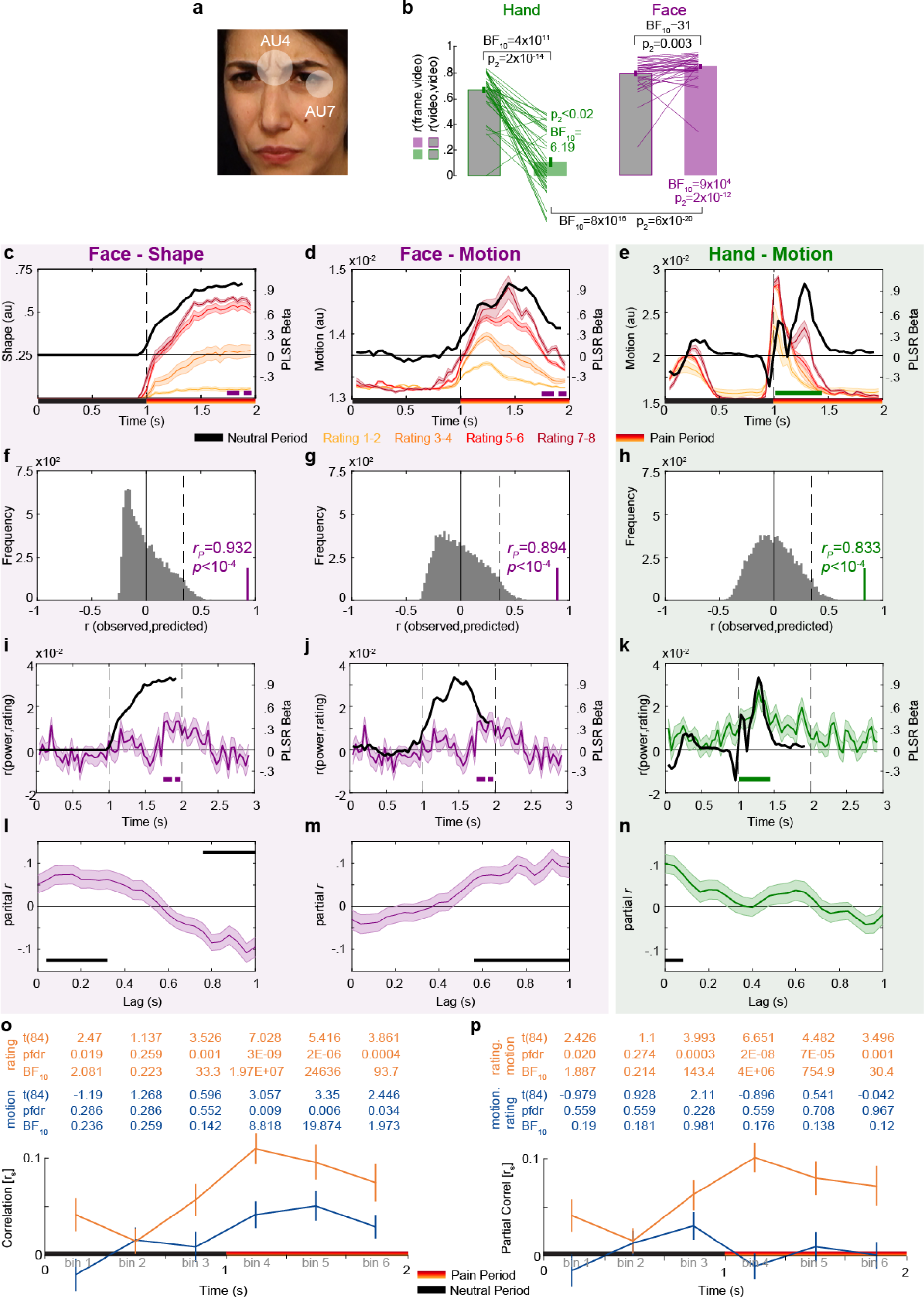
Temporal dynamics of pain rating and intensity coding in the insula broadband activity. **(a)** An example frame from the Face videos used in the online frame rating task. The average activation of the Action Units (AU) 4 and 7 was used to estimate the intensity of the shape information in Faces. **(b)** Mean±SEM correlation coefficients between each participant’s ratings in the online frame rating task and the average ratings of the other participants in the online video rating task (*r*_*S*_(frame,average_video), green and purple) compared against that between participant’s ratings in the online video rating task and the average ratings of the other participants in the same task (*r*_*S*_(video,average_video), gray) for Hands and Faces separately. Black statistics above the bars compare the respective frame and video ratings, the colored statistics compare the frame ratings against zero. The black statistic under the bars compare the frame ratings between Hands and Faces. **(c-e)** Motion and shape signals as a function of time and perceived intensity for the Face and Hand videos rated as 1-2, 3-4, 5-6, and 7-8 separately. Each colored curve represents the Mean±SEM for each rating. Purple and green bars indicate the periods with significant BBP-rating correlations for Faces and Hands, respectively. Black lines represent the partial least square regression (PLSR) beta coefficients predicting perceived intensity ratings using motion (for Hand and Faces) or shape information (for Faces). **(f-h)** Accuracy with which the motion or shape signal across all frames can be used to predict the intensity rating of the movie. The histogram shows the actual predictive accuracy averaged over cross-validation folds (green and purple) relative to the null-distribution of shuffled ratings (gray), with the median and top 5% of the null-distribution shown as full and dashed line. In all cases, the actual accuracy was higher than all 10,000 shuffling values, as indicated by *p*<10^−4^. **(i-k)** Mean±SEM time courses of the correlations between BBP and pain ratings (green and purple, as in Fig. 3g) superimposed with black lines in (c-e) for visualization of the temporal similarity between the two curves. **(l-n)** Mean±SEM lagged correlation (left) and partial correlation coefficients (middle and right) between the temporal profile of BBB-rating correlations and that of the PLSR beta coefficients for the corresponding stimulus information. For partial correlation analyses, middle panel shows *r*_*P*_(BBP(t),Motion(t+lag)|Shape(t+lag)) and the right panel shows *r*_*P*_(BBP(t),Shape(t+lag)|Motion(t+lag)). All correlations are shown for lags from 0-1000 ms in steps of 40 ms. The correlation was calculated separately for each of the 85 bipolar recordings. The Black bars represent periods of significant correlations, tested using a t-test of the 85 correlation values against zero followed by FDR correction at q=0.05. **(o)** Mean±SEM *r*_*s*_ between motion energy and BBP (blue) or between subjective rating and BBP (orange) for the 6 consecutive bins of 333 ms during the movie. All statistics are two-tailed parametric t-tests against zero, because *r*_*s*_ values were normally distributed (all Shapiro-Wilk *p*>0.05). Values are indicated in the table above each panel for each time-bin of ⅓ s. FDR correction is over the 6 bins. No *r*_*s*_-to-*z* transform was used because the *r*_*s*_ values were in the range -0.5<*r*_*S*_<0.5 for which *r* and *z* values are extremely similar. **(p)** As in (o), but partial correlations: *r*_*S*_(BBP,motion|rating) in blue and *r*_*S*_(BBP,rating|motion) in orange.

We collected data from a healthy sample of 40 participants in an online frame rating task to assess whether participants can recognize pain intensity from static frames taken at the key moment of the Face and Hand videos. Figure 4b shows that the rating of single frames of Faces were even slightly more consistent than the ratings of the entire videos from which they were taken (i.e., *r*_*S*_(frame_i_,AV)>*r*_*S*_(movie_i_,AV), *W*=193, *p*_*2*_=0.003, BF_10_=31.293). In contrast, for Hands, the rating of the frames was poor compared to the rating of the movies (i.e., *r*_*S*_(frame_i_,AV)<*r*_*S*_(movie_i_,AV), *t*_*(38)*_=11.959, *p*_*2*_=2×10^−14^, BF_10_=4×10^11^). Directly comparing the change of performance across the two stimulus types as an interaction in a effector (Hand vs Face) x stimulus (Movie vs Frame) ANOVA revealed a highly significant effect (*F*_*(1,38)*_=178.983, *p*=6×10^−16^, BF_incl_=∞). Finally, because for Hands, the accuracy was low, we also tested if the accuracy was above zero, and it was (*W*=580, *p*_*2*_=0.022, BF_10_=6.189). Hence, for Faces, static shape information was sufficient to explain the rating of the videos, while for Hands, the shape information in the frames we selected was not sufficient. It should be noted that, in principle, information contained in other frames may have contained useful information, but informal reports of some participants confirmed that they paid attention more to kinematic than configurational cues.

Leveraging the high temporal resolution of our iEEG recordings, we next asked whether the motion or the shape information better matches the timing of our intensity coding for Faces in the insula. Figure 4c shows shape information increases towards the end of the movies with rating intensity. Comparing the timing of intensity coding for the Face in the insula BBP (purple bar in Fig. 3g and 4c) with the timing of the shape information for Faces (separation between the curves in Fig. 4c) shows a nice correspondence, with both highest late in the movie. Furthemore, a partial least squares regression (PLSR) analysis indicated that the time course of shape information could predict the rating of our patients with very high accuracy (Fig. 4f). Regarding kinematics, we calculated the changes in pixel-values across consecutive frames to track the timing of motion (Fig. 4d), and this information could also predict the rating of our patients with high accuracy for Faces (Fig. 4g). Comparing the timing of intensity coding in the insula for Faces (purple bar in Fig. 4d) with the timing of motion information (separation between the curves in Fig. 4d) shows that intensity coding maximizes when motion information has already declined significantly.

We complemented these observations with a quantitative approach that estimates how the neural intensity coding lags behind the shape or motion information (Fig. 4i,j,l,m). If motion were, for instance, the driver of neural response to Face stimuli, we would expect that when motion information increases, neural responses should increase within ∼200 ms, given typical latencies for static facial expressions (Chen et al., 2009; Krolak-Salmon et al., 2003). Thus, we conducted correlation analyses to test how much the temporal profiles of the shape or the motion information are associated with the temporal profiles of intensity coding at various lags. Note that to directly contrast the predictive power of the shape and motion information, we used partial correlations. For Faces, partial correlations were positive for shape from 40-320 ms and for motion from 560-1000 ms (Fig. 4l,m). Hence, intensity coding for Faces in the insula could be driven by shape information with latencies in line with those reported in the literature for other facial expressions (Chen et al., 2009; Krolak-Salmon et al., 2003). The significant lags we revealed for the motion information are so long that motion is unlikely to have dominated the neural signals for Faces.

### 2.5. Rating-related Motion Information Could Drive Hand Intensity Coding

Motion energy is also a reliable predictor of pain intensity ratings for Hands (Fig. 4h). In our Hand videos, motion occurs at two time points: early, when the belt is lifted up, and then again, while the belt hits the hand (Fig. 4e). Pain, however, occurs only at the second point in time. This allows us to explore whether the insula is coding movement in general, or movement that is associated with pain more specifically. We thus divided the two seconds of the Hand movies in 6 segments, and asked, for each segment, how well BBP relates to motion energy in the same segment (Fig. 4o,p). Over the 85 channels, we had evidence of absence for a relationship during the neutral period that contained the period during which the belt was seen to move upwards (all BF_10_<1/3), and evidence for a relationship during the first 666 ms of the pain period when the belt is seen to slap the hand (both *p*_*unc*_<0.003, or *p*_*bonf*_<0.018 corrected for 6 bins, both BF_10_>8.8). Indeed a repeated measures ANOVA comparing the correlation values across the 6 bins confirms that the relationship between motion energy and BBP changes as a function of time (*F*_*(5,420)*_=2.9, *p*=0.014, BF_incl_=1.52). This shows that the BBP response in the insula does not code motion in general, but motion at a time when it is relevant, here to assess the pain intensity. Next, we asked whether subjective rating or motion energy was the best predictor of BBP across the 6 bins (Fig. 4o,p). Rating per se was an even better predictor of BBP than motion energy (rmANOVA, 2 predictor x 6 bin, main effect of predictor: *F*_*(1,84)*_=23, *p*=7×10^−6^, BF_incl_=13473). Interestingly, using partial correlations, we see that the correlation between rating and BBP remains highly significant when seeing the belt hit the hand even after removing what can be explained by motion energy, but we have evidence for the absence of a correlation between motion energy and BBP if removing the variance explained by rating (Fig. 4p). Together, this data supports the idea that the insula could employ motion to encode the painfulness in our Hand videos, but does not respond to simply seeing the motion of the belt, and that subjective rating of intensity appears to mediate the relationship between motion and insular response.

We also conducted the same lagged correlation analysis for Hands, as described for Faces above; that is, calculating the correlation coefficients between the temporal profile of the motion information and the temporal profile of intensity coding for Hands at various lags (Fig. 4k,n). This analysis showed that intensity coding in the insula lags behind the motion information in Hands by 0-80 ms (Fig. 4n), corroborating the above findings indicating intensity coding for Hands in the insula could indeed be driven by the motion information.

### 2.6. The Insula Contains a Surprising Number of Stimulus-specific Intensity Coding Locations

We next focused on how individual recording sites in the insula reflected perceived intensity. In the Early Period, for Hands, 21/85 (25%) showed significant intensity coding (rating-BBP correlations, n=60 trials, all *r*_*S(58)*_>0.219, *p*_*1*_<0.046), which was above chance (Binomial, 21/85 at alpha=0.05, *p*_*1*_=9×10^−10^, BF_+0_=2×10^7^). In contrast, for Faces, only 3/85 (4%) showed intensity coding in the Early Period, which is expected by chance (Binomial *p*_*1*_=0.804, BF_+0_=0.025). During the Late Period, above chance numbers of recordings showed intensity coding for Hands (14/85, 17%, *p*_*1*_=8×10^−5^, BF_+0_=201.41), and the same was true for Faces (15/85, 18%, Binomial *p*_*1*_=2×10^−5^, BF_+0_=808.49; Fig. 5a).

**Figure 5.**
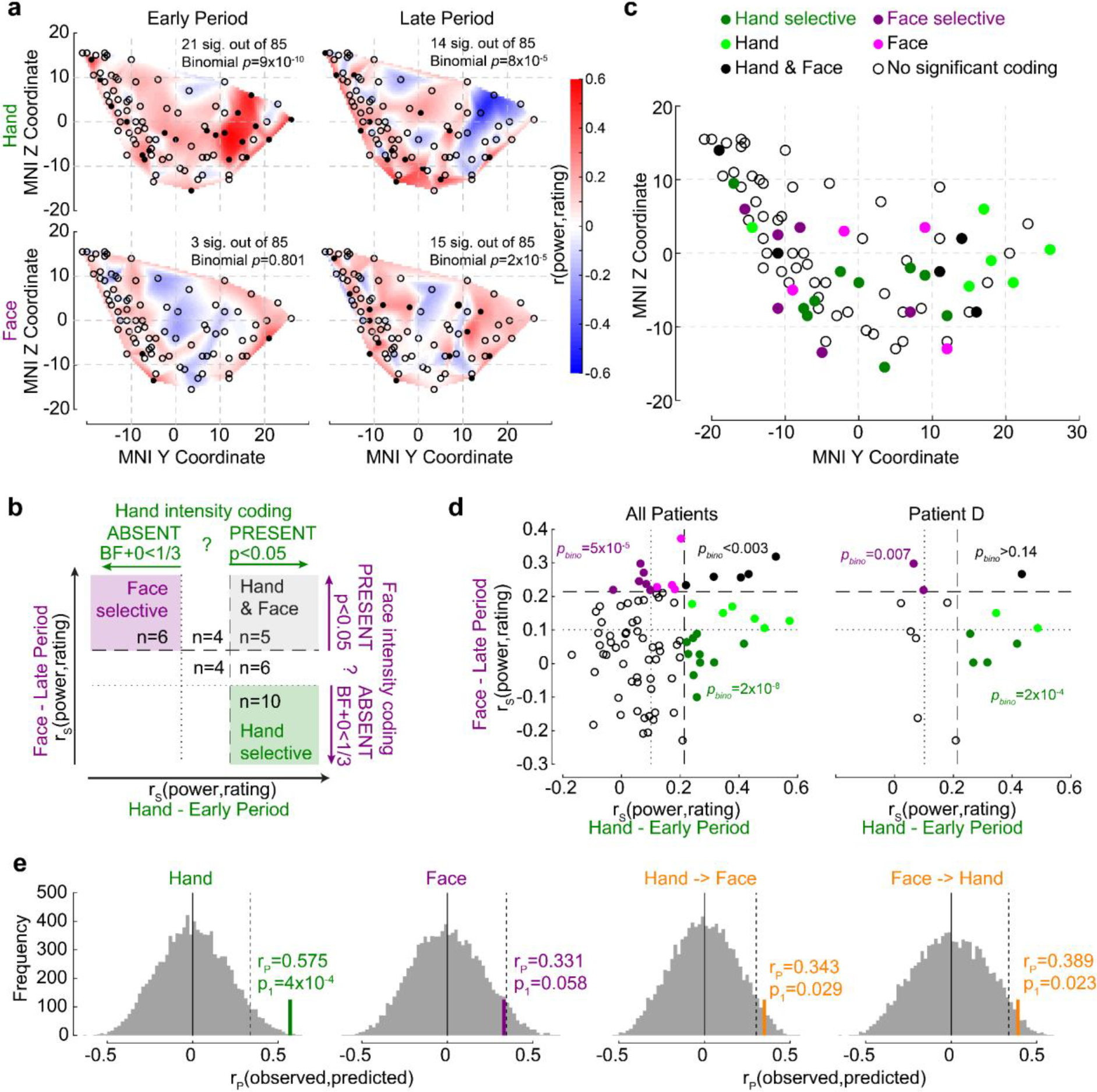
The relationship between Hand and Face intensity coding in the insula broadband activity. **(a)** Topographical maps of BBP-rating correlation coefficients for Hands and Faces in the early and late periods. Each circle is one of the recording sites (as in Fig. 1c), with filled circles indicating locations with significant correlation coefficients (*p*_*1*_<0.05). **(b)** Classification of recording locations based on their Hand (early period) and Face (late period) intensity coding. Bipolar recordings in the gray zone (n=5) significantly co-represent intensity for Hands and Faces (both *p*_*1*_<0.05, i.e., beyond dashed line). Recordings in the purple (n=6) and in the green (n=10) zone represent intensity coding selective for Faces or Hands, respectively (i.e., *p*_*1*_<0.05 for Hands and BF_+0_<⅓ for Faces, and vice versa). **(c)** Location of all 85 bipolar recordings, color-coded by selectivity as described in (b). Note that locations Hand and Face without further specification are those with *r*_*S*_ values for at least one of the stimulus types falling between the dashed and dotted lines, thus providing inconclusive evidence and showing neither significant dual coding, nor evidence of absence. **(d)** Correlation coefficients for Hands and Faces separated by coding characteristics in (b) for all patients together (left) and for an exemplary patient (right). *p*_*bino*_ refers to the likelihood to find the observed number of locations in that quadrant using a binomial distribution as detailed in Methods Section 4.1.5.8. **(e)** The left two panels depict the average correlation coefficients, together with corresponding resampling null distributions, as a measure of the accuracy of decoding intensity ratings using the partial least square regression (PLSR) beta coefficients of BBP in the early period for Hands and in the late period for Faces. The right panels are similar to the left panels, but show the accuracy of cross-decoding, that is, predicting Hand ratings from the Face BBP and vice versa. The dotted lines indicate 95^th^ percentiles of the resampling null distributions.

If the insula simply represents salience, one might expect a tight association between intensity coding for Hands and Faces, and an above chance number of locations showing dual intensity coding for both Faces and Hands. In contrast, if the insula also represents more specific information, we would expect above-chance numbers of locations with intensity coding for Faces, but not Hands and vice versa. Statistically, we infer the presence of intensity coding based on *r*_*S*_>0, *p*_*1*_<0.05, like elsewhere in the manuscript, and its absence using Bayesian statistics (Keysers et al., 2020), with BF_+0_<⅓. Plotting each bipolar recording’s *r*_*S*_ values on an x-y plot, with x representing r_S_ for Hands and y for Faces, with dashed and dotted lines at critical *r*_*S*_ values corresponding to p_1_<0.05 and BF_+0_<⅓, we define 9 quadrants, three of which are of conceptual importance (Fig. 5b): those of locations with dual intensity coding (i.e., p_1_<0.05 for Faces and Hands), those with Face-selective intensity coding (i.e., *p*_*1*_<0.05 for Faces, but BF_+0_<⅓ for Hands) and those for Hand-selective intensity coding (i.e., *p*_*1*_<0.05 for Hands, but BF_+0_<⅓ for Faces). We then used binomial tests to compare the proportion of locations falling in these three quadrants against chance, and found that all three quadrants contain more locations than expected by chance (Fig. 5d). Indeed even within a single patient, amongst simultaneously recorded channels, we find above chance numbers of Face and Hand-selective channels (Fig. 5d). Also, calculating the association between intensity coding across Hand and Face through a simple correlation of the respective r values, confirms the presence of a significant but weak, and barely worth mentioning (in a Bayesian sense) association (*r*_*K*_=0.131, *p*_*1*_=0.038, BF_+0_=1.27). Together, this shows the insula is a patchwork, with some locations representing the Hand but not the Face, others the Face but not the Hand, and a small number finally representing both in terms of intensity coding. The spatial distribution of these locations is shown in Fig. 5c.

In addition, we used a multivariate partial least square regression (PLSR) approach to assess how well the pattern of BBP across the insula can predict participants’ pain ratings. BBP across the 85 sites in the Early Period for Hands can be used to predict the patients’ average rating of the stimulus with reasonably high accuracy (n=10 trials since 1/3 of the 30 unique videos were used for testing decoding performance for each randomization, *r*_*P(8)*_=0.575, *p*_*1*_=9×10^−4^ based on reshuffled distribution), and BBP in the Late Period for Faces with almost significant accuracy (*r*_*P(8)*_=0.331, *p*_*1*_=0.058, Fig. 5e). A direct comparison of the performance of the two PLSR indicates that the performance was higher for Hands than Faces (non-parametric test across the decoding performances, *W*=944605, *p*_*2*_=9×10^−260^, BF_10_=7×10^39^). To test if intensity was encoded through similar patterns for the two stimulus types, we repeated the analyses training the PLSR on one stimulus type and testing it on the other. We found above-chance cross-decoding in both cases (Hand->Face: *r*_*P(8)*_=0.343, *p*_*1*_=0.029; Face->Hand: *r*_*P*_*(8)*=0.389, *p*_*1*_=0.023; Fig. 5e). However, when the 5 contacts that significantly co-represented pain intensity for both Hands and Faces (black dots in Fig. 5c) were excluded from the analyses, the cross-decoding accuracy fell to insignificant levels (Hand->Face: *r*_*P(8)*_=0.175, *p*_*1*_=0.153; Face->Hand: *r*_*P(8)*_=0.185, *p*_*1*_=0.149). These findings corroborate the above results, indicating that perceived pain intensity is reflected in the insula as a mixture of hand-specific, face-specific, and hand-face common representations.

### 2.7. Intensity Coding for Hands Increases Anteriorly as in a Similar fMRI Experiment

To examine the spatial distribution of intensity coding, we examined the relationship between MNI coordinates of the bipolar recordings and intensity coding (i.e., *r*_*S*_(BBP,rating), Fig. 6a). The only significant association was that more anterior recordings (i.e., more positive y-coordinates) have higher Hand intensity coding. Interestingly, we found evidence against a right-left difference (i.e., BF_10_<⅓ for x-coordinates) for the Face and Hand, providing moderate evidence against a left-right lateralization. To exclude that this finding could be driven by differences across patients, we also performed a random intercept mixed linear model using x, y and z coordinates as predictors of Hand intensity coding (without interactions) with patients as random nesting variables. This analysis confirmed the y coordinates predict intensity coding for Hands (X: *F*_*(1,79*.*23)*_=0.022, *p*_*2*_=0.881; Y: *F*_*(1,80*.*97)*_=13.23, *p*_*2*_=0.0005; Z: *F*_*(1,73*.*95)*_=0.166, *p*_*2*_=0.685).

**Figure 6.**
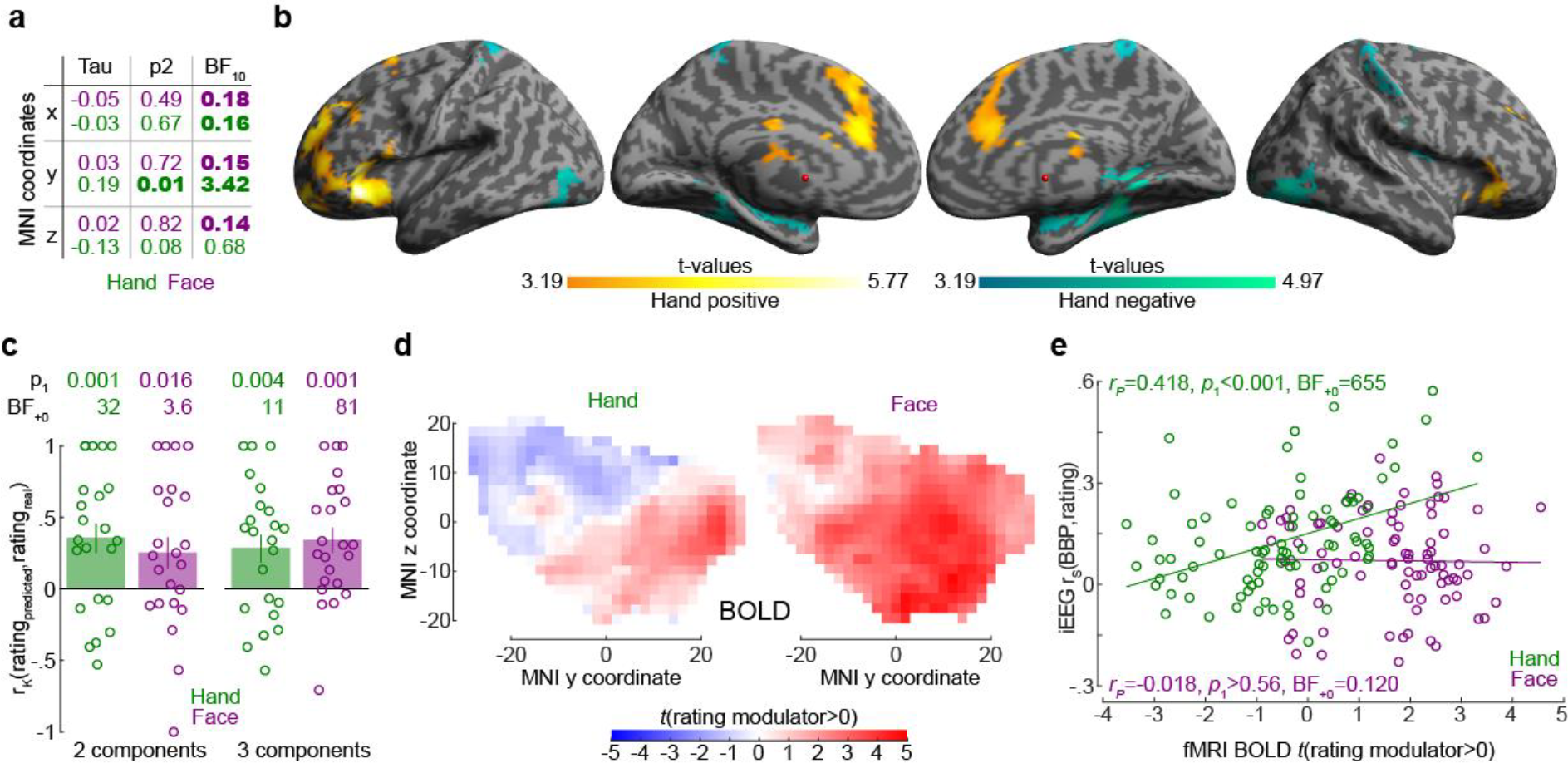
The relationship between the insula broadband and BOLD activity during pain intensity ratings. **(a)** Correlations (*r*_*K*_) between MNI coordinates and BBP intensity coding, separately for Hands (green) and Faces (purple). Bold numbers mark evidence for (BF_10_>3) or against (BF_10_<1/3) a significant correlation. Statistical values were obtained by correlating separately the x, y or z coordinate of each bipolar recording with the *r*_*S*_(BBP,rating) of each recording over all 85 recordings. Tau refers to Kendall’s Tau, *p*_*2*_ and BF_10_ the two-tailed probability and BF based on H_0_:Tau=0. **(b)** Results of the regression analysis between resting state connectivity and intensity coding for the 85 bipolar recording coordinates for Hands. Warm colors indicate significant positive, and cold, negative regression values. Results are corrected at the cluster-level at p_FWE_<0.05 using initial cluster-cutting at *p*_*unc*_<0.001, *t*(82)=3.19, and then setting the minimum cluster-size to FWEc=772 as determined by the random field theoretical calculation in SPM. **(c)** Mean±SEM of the predictive performance of a PLSR trained to predict ratings based on the pattern of BOLD activity across all voxels in the insula for different ratings. A leave-one-out cross-validation was used, and each circle represents the *r*_*K*_ between the predicted and actual rating for each left out participant, and the *p*_*1*_ and BF_+0_ values then test these n=23 correlation values against zero using a non-parametric test. Results are shown separately for Hand and Face trials and using 2 or 3 PLSR components separately. **(d)** Topography of intensity coding for the Hand (left) and Face (right), as assessed at the group level, by the parametric modulator capturing changes in BOLD activity that correlate with trial-by-trial differences in participant’s ratings. T values testing the parametric modulator >0 at the group level are shown as a function of y and z coordinate in the insula mask. For each coordinate, the maximum value across all x-coordinates within the two insulae is indicated. **(e)** Correlation (*r*_*P*_ because of normality) between the t value of the parametric modulator for the rating in the fMRI BOLD responses (x-axis) and the BBP intensity coding (computed in the early period for the Hand, green; late period for the Face, purple) in the iEEG signal (y-axis) for each of the 85 contact locations. Note that for the fMRI signal, the value is taken from the voxel closest to the MNI coordinates of the corresponding contact in the iEEG signal.

To better understand the origin of the anterior gradient for intensity coding for Hands, we performed a regression analysis between intensity coding of the 85 insular recording locations (for Hands and Faces separately), and resting state connectivity seeded at corresponding MNI locations in Neurosynth. Insular locations with higher Hand intensity coding had higher resting state connectivity with the left anterior insula and ventral prefrontal cortex (including BA44/45, OP8/9, Fp1), with the right frontal orbital cortex; with the bilateral cingulate (incl. BA24/33); and the right cerebellum (Crus I and lobules VI, VII and VIII, Fig. 6b). In line with the lack of spatial gradients for Faces in the insula of our patients, examining which voxels had higher resting state connectivity with insular locations with higher Face intensity coding did not yield any significant voxels (all *p*_*unc*_>0.001).

Finally, to compare the spatial gradient we find using iEEG with that using fMRI, we leveraged existing data from an unpublished study in our lab using a similar design to measure brain activity using fMRI in healthy participants. BOLD activity in the insula also contained significant information about the perceived intensity for Hands and Faces (Fig. 6c), and performance did not differ across Hands and Faces (t-test comparing the leave-one-subject out performance for Hands and Faces, with 2 components, *t*_*(22)*_=0.675, *p*=0.5, BF_10_=0.27; 3 components: *t*_*(22)*_=-0.39, *p*=0.7, BF_10_=0.23). For both Hands and Faces, we found a gradient along the y axis with more anterior locations showing a stronger, and more positive association between BOLD activity and rating (Fig. 6d). For Hands, across our 85 bipolar recordings in the patients, locations with higher BBP intensity coding in iEEG also show higher t values in the BOLD signal (Fig. 6e). For Faces, on the other hand, we found evidence of absence for an association of the two measures (Fig. 6e).

### 2.8. The Insula Contains Neurons with Intensity Coding for Hands and/or Faces

The insula thus displays intensity coding in a broad frequency range, including locations with Hand- or Face-specific intensity coding, as well as locations showing intensity coding for both stimulus types. To explore this representation at the level of single neurons, we analyzed the microelectrode data from the 3 patients (patients C, D, and E) that had microwires in the ventral anterior insula (pluses in Fig. 1c). Spike sorting resulted in a total of 28 candidate neurons. From these, 13 showed more spikes during the pain period than the pre-stimulus baseline. Amongst those, 8 show intensity coding for Faces and/or Hands (Fig. 7), with significant Kendall’s Tau correlations between perceived intensity (1-2, 3-4, 5-6, 7-8) and spike count during the pain period (1–2 s post-stimulus onset) for at least one stimulus type: 4/8 for Faces and 5/8 for Hands (Binomial test, Face: *p*_*1*_=0.003, BF_+0_=27; Hands: *p*_*1*_=3×10^−4^, BF_+0_=282). Considering the p_1_-value for the intensity coding, two cells (a,b) showed intensity coding for both Hands and Faces, 3 (c-e) only for Hands and 3 (f-h) only for Faces. If we additionally consider the BF_+0_ values below ⅓ as evidence for the absence of coding in the other stimulus type, we find 3 Hand-specific cells (c,d,e) and 2 Face-specific cells (g,h). Importantly, within patient D, we observe the co-existence of Hand-selective (c,d) and Face-selective intensity coding (g).

**Figure 7.**
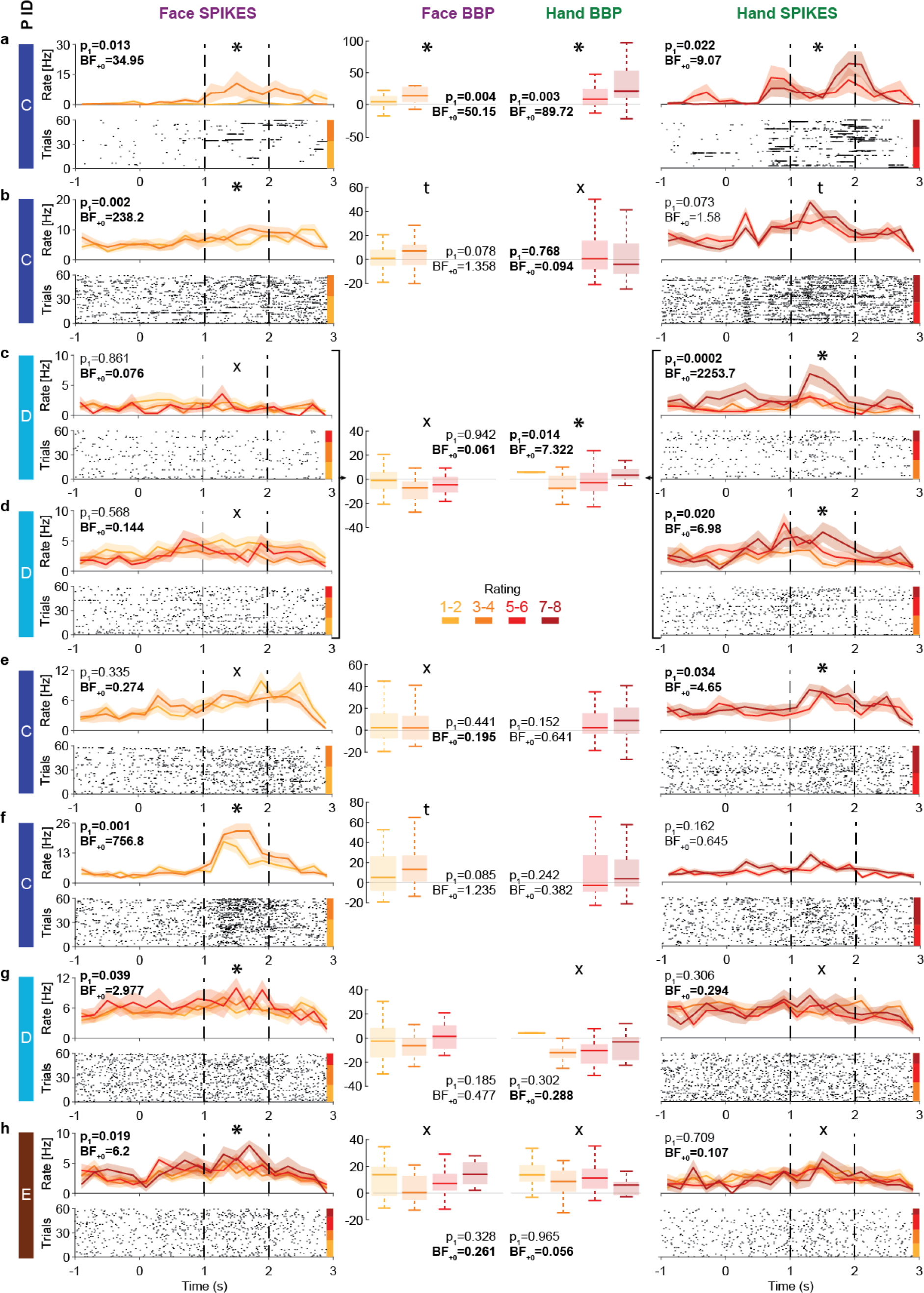
Intensity coding in the insula single-units and the corresponding broadband activity. **(a-h)** Left (Face) and right (Hand) columns display, for each single-unit, the rastergrams and peri-stimulus time histograms (PSTH) for 8 cells that showed intensity coding for at least one stimulus type. For the PSTH, each curve represents the Mean±SEM of the firing rate in each bin for trials with the corresponding rating. Not all patients gave all possible ratings in each condition. For the rastergram, trials are sorted in order of rating, with the highest ratings upmost. The colorbar next to the rastergram indicates the rows corresponding to each rating. *p*_*1*_ and BF_+0_ values result from a one-tailed test of the Kandell’s Tau between rating and spike-count in the pain period (marked by the dashed lines). *: significant intensity coding (*p*_*1*_<0.05), t: trend (*p*_*1*_<0.1), X: evidence of absence for a positive intensity coding (BF_+0_<1/3). The x-axis (time) is relative to the movie onset. The color bar on the leftmost side indicates from which patient the data is taken. Middle columns show the BBP averaged over the pain period for the microelectrode, from which the corresponding single-unit was extracted, as a function of rating as a boxplot showing the variance across trials. The box and whiskers represent the quartiles across trials, and the *p*_*1*_ and BF_+0_, the Kendall’s Tau test of the association of rating and BBP. Note that cells c and d were taken from the same microwire, and therefore have only one BBP graph.

To explore how spiking relates to BBP, we analysed the BBP from the 10 microelectrodes that yielded the 13 cells showing stimulus triggered responses. Using Kendall’s Tau correlations between BBP (20-190 Hz) and the patient’s intensity ratings (1-2, 3-4, 5-6, 7-8), and comparing these results with the coding of the cells on the same wire reveals a relationship between the two. For Hands, 2/3 microelectrodes that yielded cells with intensity coding also showed significant association between ratings and BBP (Fig. 7a,c,d). Indeed, intensity coding (i.e., correlation between intensity rating and spiking/BBP) were significantly correlated across the 10 microwires (*r*_*K*_=0.57, *p*_*1*_=0.012, BF_+0_=7.69). For Faces, only 1/5 microelectrodes with spike intensity coding cells showed significant intensity coding in the BBP, and 2/5 showed a trend. Across the wires, there was a trend towards an association between the intensity coding in the spikes and BBP (*r*_*K*_=0.34, *p*_*1*_=0.088, BF_+0_=1.63).

## 3. Discussion

Here we characterize how the insula’s iEEG activity encodes the intensity of other people’s emotions, using pain as an important category. LFPs indicate that neural activity in the insula within a broad range of frequencies, including the conventional theta, beta and gamma frequency bands, scales with the perceived intensity of pain expressed by others. Interestingly, the insula only appeared to be recruited once the perceived pain level was at least moderate: activity was increased for moderate (5-6) compared to mild (3-4) and for severe (7-8) compared to moderate. However, activity for mild pain (3-4) was not significantly increased compared to minimal pain (1-2), or baseline activity. This echoes a recent finding that BBP activity in the insula is selectively increased only once thermal stimulation is consistently painful (Liberati et al., 2020). Furthermore, we isolate a small number of insular neurons increasing their firing with increases in the intensity of pain experienced by others.

As mentioned in the introduction, BOLD signals can dissociate from neural spiking (Boynton, 2011; Maier et al., 2008). Just as V1 BOLD signals fluctuate with perception while spiking in V1 does not (Maier et al., 2008), the observation that BOLD signals in the insula fluctuate with perceived pain intensity alone cannot guarantee that neuronal spiking in the insula does. The insula’s BOLD signal could instead fluctuate with perceived intensity simply as a result of feedback synaptic input from other brain regions that encode perceived intensity (e.g., area 24 in the cingulate gyrus; Carrillo et al., 2019). The foremost impact of our broadband gamma and spiking data is thus to provide what is arguably the first evidence that the intensity of other people’s pain is indeed locally encoded in the activity of neurons in the insula.

The human insula has been in the focus of pain neuroscience as part of the pain matrix recruited by first-hand experience of pain (Ingvar, 1999). In this tradition, neuroimaging evidence for activation of the insula while witnessing pain experienced by others has led many to suggest it may represent a neural basis for empathy for pain (Bernhardt and Singer, 2012; Jauniaux et al., 2019; Lamm et al., 2011; Timmers et al., 2018). However, the insula is also recruited by a variety of tasks beyond nociception and pain empathy, including other affective states, sensorimotor functions, and decision-making under uncertainty (Craig, 2002, 2009; Uddin, 2015; Uddin et al., 2017). With so many tasks recruiting the insula, the idea has become popular that it may play a key role in attaching salience to behaviorally important stimuli (Legrain et al., 2011; Uddin, 2015; Uddin et al., 2017; Zaki et al., 2016). In the present study, we do not intend to (and cannot) address the selectivity of the insula for the pain of others over other emotions, and we do not claim the neural responses we report are pain-specific. Instead we characterize how the insula’s iEEG activity encodes the intensity of other people’s emotions, using pain as an important category, and use the agnostic terminology of ‘intensity coding’, rather than ‘pain intensity coding’ throughout the paper. That the insula’s broadband activity correlated with motion energy in our Hand stimuli only while the motion was associated with pain (during the slapping), but not while motion was innocuous (during the initial belt-lifting), shows that the effects we measured cannot be reduced to visual motion detection. That perceived pain intensity was mediating the association between motion and broadband activity further speaks against an interpretation of our results as reflecting unspecific motion processing. Future experiments using a wider gamut of control stimuli, that are matched in motion but differ in emotions will be critical to address the specificity of the responses we describe, and hence, what state they could reliably signal (Zaki et al., 2016). For the Hand stimuli, this could include seeing hands experience a range of different salient affective experiences, such as a hand being hurt, being caressed and being rejected in addition to a neutral hand-shake (Meffert et al., 2013). For the face, this could include disgusted, angry, fearful, and happy facial expressions matched for motion.

An important, and somewhat related, question has been whether pain or salience cues are represented in a modality-specific or modality-general manner in the insula. fMRI studies have shown that the anterior insula is coactivated with different brain regions depending on whether the pain of others is conveyed via indirect cues or via the actual body part, such as the hand, that directly receives the noxious stimulation (Gallo et al., 2018; Jauniaux et al., 2019; Keysers et al., 2010; Lamm et al., 2011; Timmers et al., 2018). Here, we provide electrophysiological measures of neural activity that speak to that issue. We focus on the broad-band gamma signal known to have comparatively high spatial specificity and be closest to neural spiking (Bartoli et al., 2019; Buzsáki et al., 2012; Miller et al., 2014), and find a mixed organization, consisting of modality-specific and -general locations in a partially intermixed layout. That is, we found locations with broadband activity and spiking associated with perceived intensity for the Hand, but not the Face; others associated with the Face, but not the Hand; and others still associated with perceived intensity for both.

Leveraging our high temporal and spatial resolution, we found that locations that showed intensity-coding for the Hand stimuli have activity timing echoing the timing of pain-related motion cues with relatively short latencies <100 ms, and that the association of motion energy and broadband activity is mediated by the perceived intensity. Locations that show intensity coding for the Face appear to have activity echoing the timing of shape information with latencies no longer than 320 ms. These latencies are in a range similar to that found following nociceptive stimulation (Liberati et al., 2020) or static disgusted facial expressions (Chen et al., 2009; Krolak-Salmon et al., 2003). Using automated software to detect the level of activation of the facial action units 4 and 7 (i.e., lowering the eye-brows and tightening the eye-lids), we found that this information suffices to predict participants’ rating of the stimuli with high accuracy, and followed the time course of the neural activity in the Face intensity encoding locations well enough to suggest that it provides traction on the analyses of dynamic pain-related facial expressions.

An important consideration is whether this selectivity for Face or Hand stimuli could simply originate from some participants finding our facial stimuli more salient, and others the hand stimuli, particularly, given that our patients rated the Hand stimuli with slightly higher pain intensity than our Face stimuli. That we find Face and Hand selectivity side-by-side in simultaneously recorded locations and neurons in single patients suggests that this cannot suffice to explain our data. This is because if a patient were to find Hand stimuli more salient than Face stimuli, and the insula simply codes saliency, we would expect to find Hand-specific intensity coding in that patient’s insula, but we wouldn’t expect to find side by side locations with Hand-specific and Face-specific coding. Our findings instead suggest that different locations might encode behaviorally relevant (and hence salient) information about the intensity of the pain of others with some degree of specialization for a particular type of information (Hand vs Face). The insula intensity coding we measure could have a dual function: help perceive the intensity of other people’s emotions and tag intense emotions as salient, thereby reconciling the notion that it contributes to empathy and to saliency. One peculiar observation in our data is that broadband power for the Hand stimuli did not increase monotonically as a function of reported pain intensity, but in a J shape, with higher power for the lowest than second lowest rating. Some have argued that the insula may be part of an active inference circuitry in the brain that learns to predict emotionally relevant states and stimuli (Seth and Friston, 2016). In such a perspective, a J shaped response curve could reflect a combination of underlying neurons representing the intensity of other people’s emotions (with a monotonic increase in activity) and others representing the prediction error (which for randomly presented intensities would have a U shape centered on the average intensity). Future experiments could contrast responses to a given pain intensity in blocks of high and blocks of low average presented intensity to disentangle the effect of observed and expected intensity on neural responses in the insula and shed further light on this predictive component.

On the other hand, we also found evidence that some locations and cells represent intensity coding for both the Hand and the Face stimuli. In addition, if we train a partial least square regression to predict perceived intensity from the activity pattern across all recorded locations, we find that training the decoder on Hand activity pattern and testing it on Face activity patterns (or vice versa) leads to above chance decoding. This confirms that the insular representation of intensity can support stimulus independent decoding - despite our partial least square regression not being biased to focus on signals that do generalize well across stimulus types. This provides an electrophysiological basis for recent fMRI studies that show that stimuli depicting situations in which others’ experience of pain or not can be discriminated using the same pattern across hand and face stimuli (Zhou et al., 2020).

In addition to the broadband results we report in detail, we find that theta power increases with perceived intensity. Given a growing animal literature establishing that interareal theta synchronization promotes learning about threats (Likhtik et al., 2014; Likhtik and Gordon, 2014; Taub et al., 2018; Tovote et al., 2015), examining the coherence in the theta range across iEEG electrodes in different brain regions during pain observation may in the future shed light on how humans learn about safety through others.

Spatially, finally, we found that Hand intensity coding was enriched in the anterior dorsal insula, where we also found the largest proportion of locations encoding both Hand and Face intensity. This anterior bias was also observed in our BOLD signal for similar hand stimuli. A recent meta-analysis identified that the most consistent BOLD activations when observing limbs in painful situations within the insula occur bilaterally around MNI coordinates y=13 and z=10 (Jauniaux et al., 2019), which closely matches where we find the highest density of Hand intensity coding (Fig. 5a). Interestingly, locations with higher Hand intensity coding have increased connectivity at rest with extra-insular regions involved in processing two relevant stimulus dimensions. Connectivity was higher with cerebellar lobules VI, VII, and VIII, and with the inferior frontal gyrus, all of which are recruited by (Abdelgabar et al., 2019; Caspers et al., 2010; Gazzola and Keysers, 2009) and necessary for (Abdelgabar et al., 2019; Keysers et al., 2018; Pobric and de C. Hamilton, 2006) perceiving the very kinematics of hand actions we find to be good predictors of BBP activity in the current study. Connectivity is also higher with the mid- and anterior cingulate cortex associated with pain witnessing in humans (Jauniaux et al., 2019; Lamm et al., 2011; Timmers et al., 2018) and that contain neurons with pain intensity coding in rats witnessing the pain of other rats (Carrillo et al., 2019).

With respect to Faces, somewhat surprisingly, we did not find a clear spatial clustering of intensity coding in our electrophysiological data. Overall, our ability to decode perceived intensity from our Face stimuli was also lower than that from the Hand stimuli. In that context, it is important to reflect on the fact that, despite our efforts to match the perceived intensity based on previous data (Gallo et al., 2018), patients (and to a lesser extent an age and gender matched control group) perceived the Hand stimuli as more intense than the Face stimuli. Given that responses were strongest for the highest rating, and that this rating was given more rarely for Faces than Hands, this difference could have contributed to making our Face results less consistent. At the same time, the variance in rating, a critical factor for determining the efficiency with which a regression can detect the presence of a rating-BBP relationship, did not differ across Hand and Face. Together, this difference in perceived intensity cautions us against overinterpreting the lack of detectability of a topographic organization of responses to Faces. Indeed, in our BOLD data, where decoding performance was similar for Hand and Face stimuli, Face intensity coding also had a clear spatial organization, being stronger more anteriorly, and meta-analyses show the left anterior insula to be reliably recruited by the observation of painful facial expressions (Jauniaux et al., 2019). However, that we found fewer locations and less reliable spatial organization for Face than Hand intensity coding does dovetail with recent meta-analyses of the fMRI literature show that, when comparing studies showing limbs in painful situations with those showing painful facial expressions, the insula is more reliably recruited by the sight of limbs (Jauniaux et al., 2019; Timmers et al., 2018). Indeed, that we find a macroscopic organization for the Hand, but not the Face, intensity coding is echoed at the mesoscale: microwires with cells with Hand intensity coding also tend to show Hand intensity coding in the BBP signal that is thought to pool the spiking of many neighbouring neurons, but the same is not true for the Face. In terms of lateralization, we find that our data is more likely if one assumes that both hemispheres have similar intensity coding, than if one hemisphere were dominant. This echoes the fact that during noxious stimulation on the right hand, both insulae show significant iEEG responses (although slightly stronger in the left insula; Liberati et al., 2020) and that fMRI fails to find robust lateralization of responses to empathy for pain (Jauniaux et al., 2019; Timmers et al., 2018).

## 4. Materials and Methods

### 4.1. iEEG Experiment

#### 4.1.1. Participants

##### 4.1.1.1. Patients

Depth electrode recordings were collected from 9 epileptic volunteers, admitted at the Amsterdam UMC to localize seizure origin. Patients received initial study information from the neurosurgeon and provided informed consent to the experimenter before the surgery occurred. Our single session experiment started on average 4 days after surgery (std = 1.89 days). Two patients were excluded from the analyses due to poor behavioral performance, and seven were included (4 females, 34.3y±9std, Table 2). The study was approved by the medical ethical committee of the Vrije University Medical Center (protocol 2016/037) and each patient signed a written informed consent according to the Declaration of Helsinki.

Clinical investigation revealed that for all our patients, the epileptic incidents did not appear to originate around the electrode contacts in the insula that we analyzed here. In addition, for four of them, recordings pointed to origins of the epilepsy to be clearly outside the insula, leading to the surgical removal of extra-insular regions (Table 2). Finally, for the remaining 3, no clear origin for the epilepsy could be localized, but there was no indication that the insula was involved in the initiation of the attacks.

##### 4.1.1.2. Control participants in the online video rating task

To assess whether the behavior of the patients was representative of the general population, we compared patients’ ratings with those of ninety-three volunteers (54 females, 32.7y±9std), who took part in an online version of the video pain rating task. The matching with the 7 patients was done by only including age- and gender-matched Dutch participants, and was successful (Table 2). The study was approved by the local ethical committee of the University of Amsterdam (2021-EXT-13608) and each participant signed an online informed consent form to participate in the study.

##### 4.1.1.3. Control participants in the online frame rating task

To determine if participants could use shape information available in single frames to determine pain intensity in Hand and/or Face stimuli, forty volunteers (23 females, 33.7y±9std) from the same group that also performed the online video pain rating task participated in the online frame rating task, so they were already familiar with the videos and had a better understanding of where the single frames came from. This also allowed us to directly compare how they rate single frames with how they rated the movies from which the frames were taken. They were selected to approximate the age and gender distribution of the patient group.

#### 4.1.2. Stimuli and Procedure

##### 4.1.2.1. Video rating task

The 2 s videos were generated as in Gallo et al. (2018) and showed a Caucasian female receiving either electrical shocks to the hand (reaction conveyed by the facial expression only; Face condition) or a slap with a belt to the hand (reaction conveyed by the hand only; Hand condition). Hence the location of the noxious stimulation was maintained across conditions (dorsum of the left hand), but the cues through which participants could deduce the painfulness differed. All videos started with 1 s of baseline: neutral facial expression for Face and static hand for Hand stimuli. Movies were cut, so that evidence of pain started at 1 s (Fig. 1a). Before the experiment, participants were instructed to rate pain intensity (“How much pain do you think the person felt?”) on a scale from 0 (no pain at all) to 10 (the worst imaginable pain). To reassure patients that no real harm was never inflicted to the actor in the movie, they were informed that during video recording stimulations in the 9-10 range were never used. Participants had to rate pain intensity after each video at their own pace, using 4 keyboards-keys (Fig. 1b). Only the relevant keys were presented on the screen, intensities were not indicated. Patients watched each of the 60 videos (30 Hand, 30 Face) twice in fully randomized fashion with a random interval of 1.5 s±0.5. The videos were matched in terms of intensity and standard deviation based on a validation in Gallo et al.(2018)

##### 4.1.2.2. Online video rating task

The stimuli and the task were the same as in the electrophysiology experiment, except each video was presented only once. Prolific Academic (https://www.prolific.co/) was used to recruit participants and the experiment was implemented on Gorilla (Anwyl-Irvine et al., 2020; https://gorilla.sc/).

##### 4.1.2.3. Online frame rating task

The task was similar to the pain rating experiment, except still frames instead of the full videos were presented for 2 sec. For faces, frames at the 1.8 s of the Face videos were used (except for one video where the eyes were closed at 1.8 s, so the frame was taken at 1.68 s). This time-point was selected because facial expressions were most pronounced towards the end of the movies, and more formal analyses confirmed that this corresponds to a time where shape information plateaued (Fig. 4c). To use a comparable stimulus set for Hands, which portrayed maximal configuration information, we selected from each Hand video separately, the frame at which the hand was maximally depressed by the force of the belt slap (time point Mean±SD=1.001±0.013 s).

#### 4.1.3. Data Acquisition

Patients were implanted with Behnke-Fried depth electrodes (Ad-Tech Medical Instrument Corporation; Fried et al., 1999) targeted at the right or left, anterior or posterior insula. Electrodes were inserted via a guide-tube under the guidance of an online stereotactic positioning system. They consisted of a silastic hollow tube with 9 to 12 platinum outer macro contacts, 1.28 mm in diameter, 1.57 mm in length with the first two macro contacts spaced 3 mm from each other and the rest spaced 5 mm from each other. This hollow tube had 9 platinum microwires (8 recording and 1 reference contact) running through it, each 38 micron in diameter, protruding as a “pigtail” formation out of the tip of the electrode. Macro contact recordings were amplified using unity gain, DC amplifiers (Braintronics BrainBox 1166 system), low-pass filtered at 1500Hz (−3dB point, -12db/octave) and sampled at 32768 Hz. The digital signal was decimated to a final sample-rate of 512 Hz or 1024 Hz and was pre-filtered with a 3 section FIR equiripple filter (0.01dB passband ripple) with the passband set to 1/3 of the sample frequency and the stopband set to 1/2 of the sample frequency. Signals from the micro contacts were amplified with respect to a skull-screw ground using a unity gain HS-9 head-stage amplifier (NeuraLynx). The signal was high-pass filtered at 1 Hz and low-pass filtered at 5 kHz and had a sampling rate of 32kHz. There were a total of 85 macro electrodes and 32 micro wires across all patients in the insula that we recorded from.

#### 4.1.4. Electrode Localization

For each patient, the T1 structural magnetic resonance (MR) image taken before the electrode implantation surgery and the computerized tomography (CT) scan taken after the electrode placement were co-registered (Fig. 1d). Using SPM12 (www.fil.ion.ucl.ac.uk) the T1 image was segmented to determine the normalization parameters, and MR and CT images were then normalized to the MNI space using these parameters. CT scan and gray matter were overlaid with insula probability maps (Faillenot et al., 2017) and macro contacts within the boundaries of the insula map were detected based on detailed visual examination using MRIcron (https://www.nitrc.org/projects/mricron). Since macro contact recordings were analyzed in a bipolar layout, the coordinates of each bipolar recording was estimated as the midpoint of its macro contacts (Supplementary File 1). The coordinates of the microwires were determined at the tip of each electrode (Supplementary File 1).

#### 4.1.5. Data Analysis

##### 4.1.5.1. General Statistical Approach

Much of the analyses in this paper assess intensity coding, which examines the relationship between brain activity (measured based on LFP, spiking or BOLD activity) and rating. Because the rating of pain intensity was along discrete categories (1-2, 3-4, 5-6, 7-8), that might be linear, but is certainly ordinal, we tend to use association measures that are appropriate for ordinal scales when we relate brain activity to the rating of a single participant. That includes Spearman r in most of our MATLAB codes, when analyses need to be repeated for every electrode because it is the most widely used rank-order correlation metric. We use Kendall’s Tau, when using Bayesian analyses implemented in JASP, because these analyses are not yet available for Spearman r. When examining the association between variables that are more continuous and normally distributed, we use Pearson’s r.

When using t-tests, we examined normality using the Shapiro-Wilk test. If normality is preserved, we report t tests and t-values; if not, we use Wilcoxon signed rank or Mann-Whitney U tests, as indicated by W or U values, respectively. When possible, or when evidence of absence is important for the interpretation of the data, we supplement the frequentist p values with Bayesian statistics calculated using JASP (https://jasp-stats.org). We use the abbreviations p_1_ to represent one-tailed p values, and p_2_ for two-tailed p values. BF_10_, and BF_01_ represent relative evidence in form of the Bayes Factor for H_1_ and H_0_, respectively, when two-tailed hypotheses are used. When we look for intensity coding, we focus here on positive intensity coding, and thus use directed hypotheses, marked with p_1_ or BF_+0_ or BF_0+_, with the + indicating a directed H_1_, using conventions as in Keysers et al. (2020). It should be noted that the use of one-tailed statistics, which is sometimes criticised when exclusively using frequentist statistics, has important advantages when combining the frequentist with a Bayesian framework, in that it increases the sensitivity for falsifying the alternative hypothesis in a Bayesian framework. Corrections for multiple comparisons are performed when repeated testing is done across time-points using either FDR corrections, or by calculating a null distribution of cluster statistics. When testing multiple neurons, or multiple bipolar recordings, we do not correct for multiple comparisons when attributing a property to a location, as this would result in changing the property of a location based on how many locations have been tested. Instead, we then examine whether the number of electrodes with a certain property exceeds the number expected by chance using binomial distributions.

##### 4.1.5.2. Behavioral Analyses

To explore whether patients were impaired in their ability to perform the task, our rationale was to consider the average rating of all control participants as the normative rating. We then compared the vector of 30 ratings (one per movie for 30 movies) of each member of the control group against the average of the other members of the control group to define a distribution of how far from the normative rating healthy volunteers tend to fall. For the patients, we compared their ratings against the average rating of the control group, and compared how similar patient ratings were to the normative average against the distribution of how similar left-out control participants are to the normative average. We calculated three metrics of similarity: the Spearman’s rank order correlation, the slope, and the intercept of a simple linear regressions between the ratings of each of the patients and the average rating of all control samples.

##### 4.1.5.2. Preprocessing of LFPs

To reduce artefacts and extract local signals, iEEG macro contact recordings were digitally re-referenced in a bipolar layout (Fig. 1d). This generated 85 bipolar recordings from 102 contacts in the insula, with patients having between 5 and 19 bipolar channels (Fig. 1c, Supplementary File 1). Re-referencing attenuated 50 Hz noise sufficiently to omit digital filters that distort data. Continuous recordings were separated into trials of 4 s: 1 s premovie baseline, 2 s video, and 1 s postmovie. Trials were visually checked for ground failure and amplitude saturation (none was detected), downsampled to 400 Hz, and detrended.

##### 4.1.5.3. Time-frequency Decomposition of LFPs

A sliding window Hanning taper based approach was used for each trial with the following parameters: frequencies from 1 to 200 Hz in steps of 1 Hz; time points from -1 s (relative to movie onset) to 3 s in steps of 0.0025 s; and for each frequency, a single sliding Hanning taper window with the duration of 8 cycles (maximum=1 s; minimum=0.1 s). Trials were expressed as % power change relative to baseline (−1 s to 0 s) separately for each frequency: y(t)=(P(t)-P_0_)/P_0_), with P_0_=average of baseline. Points with y(t) ±10 standard deviations from the mean of the other trials were excluded to not reject entire trials, but only outlier time-frequency points in some trials (rejections were rare, mean rejected time-frequency points=0.0032%±0.0035std).

##### 4.1.5.4. Intensity Coding in LFPs

In the LFP signal, we consider that a bipolar channel shows intensity coding if its trial-by-trial power variations correlate positively with the variation in the pain intensity reported by the patient. We always coded the 1-2, 3-4, 5-6, 7-8 rating options as 1, 2, 3, 4. For each bipolar recording, we then calculated the Spearman’s rank correlations (due to the ordinal nature of intensity ratings) between the patient’s intensity rating and power estimate over all trials either for each time-frequency intersection separately, or within a certain power-band. A one-sample t-test was used to test whether the average correlation over the 85 bipolar recordings differed from 0. Correlations were not Fisher r->z transformed, because r and z values remain virtually identical for -0.5<*r*<0.5, which is the range in which the correlations we examined remain. Results were cluster-corrected based on a circular shift procedure, in which the time-frequency profile of the correlation coefficients for each contact was randomly time-shifted, all such time-shifted profiles analyzed with one-sample t-tests against 0 at each time-frequency intersection separately as described above, and the sum of the largest significant positive and negative clusters were calculated separately. As described in Maris and Oostenveld (2007), this procedure was repeated for 1000 iterations, which generated maximum sum significant cluster distributions for the positive and the negative clusters expected by chance. The probabilities of the observed cluster sums were calculated under the corresponding null distributions for checking statistical significance in cluster correction.

We performed a similar analysis to identify time-frequency bands with significant intensity coding when separating the Hand and Face trials. Note that including half the number of trials makes this analysis less powerful than the Hand and Face combined analysis, which is why we used the broadband frequency band (20-190 Hz) resulting from the combined analysis. To directly compare the time-frequency profiles of intensity coding for Hand vs Face trials, we performed a similar analysis for cluster correction, except that for each time-frequency point separately, the Hand and Face correlation coefficients distributions across the 85 electrodes were directly compared with a paired-samples t-test, and, instead of the circular shift randomization procedure, trials were randomly assigned as Hand and Face trials at each step of the 1000 iterations for generating the null distributions.

##### 4.1.5.5. Resampling LFPs from the Entire Brain

To test whether the BBP-rating association observed in the insular electrodes was enriched compared to what could have been observed in any 85 bipolar recordings anywhere in the brain, we made use of not just the insular, but all the intracranial macro-electrodes that were implanted in the same patients. The recordings from these electrodes were preprocessed exactly as described for the insular electrodes. The seven patients included in these analyses had between 91 and 149 (Mean±SD=114±20) bipolar recordings distributed throughout the two hemispheres and various regions of the four brain lobes. The BBP-rating Spearman correlation coefficients of each of these electrodes were entered into a resampling method, in which, in each of the 100,000 iterations, from each patient, a random subset of these correlation coefficients were selected. The number that was selected for each patient was determined by the number of insular electrodes that patient had; that is, since Patient A had 5 insular electrodes in the main analyses, 5 random electrodes were selected from the entire brain in these analyses. This was done to ensure that possible patient-specific biases in the analysis of insular electrodes were maintained in these analyses. This way, in each iteration, a total of 85 electrodes were selected randomly from the entire brain and tested with a one-sample t-test against 0. The resulting 100,000 t-values from all the iterations were used as the null-distribution to test whether the t-value observed in the insula was greater than what would be expected if we were not focusing on the insula and were randomly sampling from the entire brain. It is important to note that while the anatomical location of electrodes in and close to the insula were carefully determined, this manual procedure was not performed for electrodes clearly outside the insula, making it possible that some of these extra-insular electrodes included in this resampling were located in the white matter or cerebrospinal fluid, and this analysis should thus be considered with a grain of salt.

##### 4.1.5.6. Extracting Shape and Motion Information from Videos

A recent systematic review has revealed that facial expressions of pain are most consistently characterized by lowering of the brow and tightening of the eye-lid (Kunz et al., 2019), corresponding to facial action units 4 and 7 (Ekman and Friesen, 1978). More specifically, research has evidenced that people fall into four clusters that differ in how they express pain (Kunz and Lautenbacher, 2014), and our protagonist fell within cluster IV, who express pain by furrowing brows and tightening eyes, but not opening the mouth or wrinkling the nose. Accordingly, we quantify the painfulness expressed in the shape of our protagonist’s face based on facial Action Units AU4 and 7. To get a replicable and objective measurement of these AU, we used the FaceReader software (Noldus, the Netherlands), which uses a deep convolutional neural net to automatically extract the level of activation of the classic facial action units (Ekman and Friesen, 1978). FaceReader reliably obtained estimates for the facial actions units 4 and 7 from all but 3 frames from our 30 movies, and we thus quantified the pain related shape information contained in each frame of our movies as the average of action units AU4 and AU7. When applied to the frames used in our psychophysical experiment mentioned above, the average activation of AU4 and AU7 correlated at r_p_=0.95 with the average rating from human observers of the same static images, validating the utility of this automated signal. Unfortunately, we found no software that could estimate muscular contraction from the hand in a similar way, and we thus did not see an obvious way to extract shape information from the Hand stimuli. Given that participants are also very poor in their ability to rate painfulness from static frames of the hand configuration in our stimuli, as shown by our psychophysics, we felt that not quantifying shape information for the Hand stimuli was acceptable.

To quantify motion over time for each video, we use motion energy, an established and objective way to extract dynamics from any movie, using the average of the Euclidean distances between the RGB vectors of the corresponding pixels across every two consecutive frames.

##### 4.1.5.7. PLSR Decoding of Intensity Coding from Shape and Motion Information

To identify when motion or shape information may contribute to predicting the overall intensity rating R of the movie i, we used partial least squares regression (PLSR) analyses using the ‘plsregress’ function in MATLAB, with a single component. For the Hand, where no shape information was available, the predictor for each movie i was the motion M at frame t, and the plsregress thus identified the weights (B), such that R(i)=M(i,t)B(t)+B_0_(i). For the Face, where both motion and shape information S was available, as the average of AU4 and 7 in each frame, we concatenated M(i,t) and S(i,t) into a single predictor X to identify weights such that: R(i)=XB+B_0_(i). We used PLSR here in particular, because both M and S have high temporal autocorrelations and are mutually correlated as well, and PLSR are well suited for such cases.

Second, we can use the PLSR method to see how accurately the motion and/or shape profile across the entire movie can be used to predict the rating of the patients, using a cross-validation. The predictive accuracy for both the motion and shape time course were calculated in 1000 iterations. In each iteration, the videos were randomly divided into three equal-sized samples. For each of these three samples separately, the remaining two samples were used to calculate PLSR beta coefficients, which were then used to predict the ratings of our patients. The Pearson correlation coefficient between the actual ratings and the predicted ratings were taken as measures of predictive accuracy and averaged across all iterations. A similar procedure was also applied to calculate the null distribution of such correlation coefficients, in which, the 1000-iteration step described above was repeated for 10000 times, each time with a different randomly shuffled version of the observed ratings. The probability of the observed decoding accuracies were then estimated by ranking the accuracy based on the actual ratings within the distribution of shuffled ratings. Figure 4f-h shows that motion and shape information each allows one to predict movie ratings with high accuracy.

##### 4.1.5.8. Probability of Face-selective, Hand-selective, and Dual Intensity Coding LFPs

Correlations between BBP and rating were thresholded as significant or as providing evidence of absence as follows (Fig. 5b). At n=60 trials, values above *r*=0.214 show a significant positive association (*p*_*1*_<0.05). Values below *r*=0.085 provide evidence for the absence of a positive association (BF_+0_<1/3). Intermediate values are inconclusive (Keysers et al., 2020). Both the frequentist and Bayesian criteria we use here are subject to type I/II errors, and we thus asked whether the number of bipolar recordings we find in these quadrants is above what we would expect by the probability of these errors. For the frequentist criterion, *p*_*1*_<0.05, we expect 5% of locations to be classified as showing significant intensity coding even if H_0_ was true (i.e., despite no real intensity coding). With regard to the dual-coding quadrant that we are interested in, two types of errors could be made. The most likely misclassification is for a location showing one intensity coding to be mistakenly classified as having dual intensity coding. To test if we have above chance numbers of dual-coding locations, we thus take all the locations with Hand intensity coding (21/85), and ask amongst those, whether finding 5 also showing Face intensity coding is more than what we expect using a binomial n=21, k_success_=5, α=0.05) and the results showed clear evidence that there are more locations also representing the Face amongst the Hand locations (*p*_*1*_=0.003, BF_+0_=17.09). The same could be done by looking whether 5 Hand intensity coding is overrepresented amongst 15 Face intensity coding locations (p_1_=0.0006, BF_+0_=117). A less likely misclassification is for a location that shows neither intensity coding to be classified as having both (=0.05^2^). Making 5 such misclassifications amongst 85 recordings is also highly unlikely (*p*_*1*_=3×10^−6^, BF_+0_=4446). For the Bayesian criterion, BF_+0_<1/3 this calculation is more difficult to perform, because Bayesian criteria are not defined directly based on a false positive rate. However, Jeffreys (1939) chose the bound of BF<1/3 as evidence of absence or presence precisely because it roughly corresponds to a p=0.05 with a standard prior on the effect sizes in H_1_. We can thus, as a reasonable approximation, assume that if H_1_ is actually true, and a location thus shows significant positive intensity coding, only 5% would be falsely classified as showing evidence against H_1_. With that approximation, amongst the 21 Hand intensity coding locations, it is highly unlikely to encounter 10/21 showing evidence that they do not encode the Face if in reality they did: (Binomial with n=21, k=10, α =0.05, *p*_*1*_= 2×10^−8^, BF_+0_=2×10^6^). Even if the the false rejection rate were much higher (e.g., α =0.25), 10/21 remain unlikely (Binomial *p*_*1*_=0.025, BF_+0_=4.2). Similarly, amongst the 15 locations with significant Face intensity coding, finding 6 with evidence for not encoding the Hand is again unlikely (Binomial n=15, k=6, α=0.05, *p*_*1*_=5×10^−5^, BF_+0_=1334). We can thus conclude that selectivity is over-represented in the insula compared to what we would expect if all neurons showing selectivity for one stimulus type would also show selectivity coding for the other.

The same analysis was applied to an exemplar participants (Fig. 5d). Using the same logic in that patient we find that the number of Hand selective locations (*p*_*1*_=2×10^−4^, BF_+0_=701) and the number of Face selective locations (*p*_*1*_=0.007, BF_+0_=37) are surprising, but the number of dual-selectivity (1/15) is not surprising amongst the 3 Face (*p*_*1*_=0.143, BF_+0_=1.9) or 7 Hand (*p*_*1*_=0.3, BF_+0_=0.48) coding locations.

##### 4.1.5.9. PLSR Decoding of Intensity from the LFP Insula Activity Pattern

To explore how well the pattern of activity across all 85 bipolar recordings reflects the perceived intensity reported by our patients we applied a PLSR regression approach similar to that used to infer how well shape or motion predicts ratings, except that instead of using motion over 50 frames, we used BBP power over 85 electrodes. Specifically, BBP across the 85 sites, averaged over the early period for Hand videos and in the late period for Face videos, were separately used as predictors in two separate PLSR analyses predicting the participants’ average pain ratings. The decoding accuracies were each calculated in 1000 iterations. In each iteration, the videos were randomly divided into three equal-sized samples. For each of these three samples separately, the remaining two samples were used as a training set to calculate PLSR beta coefficients, which were then used to predict the ratings in the remaining test sample. The Pearson correlation coefficient between the predicted ratings and the actual ratings of the patients was then taken as a measure of decoding accuracy and averaged across all iterations. We used Pearson here because the data was normally distributed, and we compared two ratings. A similar procedure was also applied to calculate the null distribution of such correlation coefficients, in which, the 1000-iteration step described above was repeated for 10000 times, each time with a different randomly shuffled version of the original ratings. The probability of the observed decoding accuracies were estimated under the corresponding null distributions as the rank of the actual average accuracy against the shuffled accuracies. We first performed this analysis within each stimulus type - i.e., we trained and tested on Hand stimuli or we trained and tested on Face stimuli. Then, to explore whether the pattern of activity could generalize across stimulus type, we also performed cross-decoding analyses where we trained on one stimulus type (e.g., we determined the PLSR weights using ⅔ of Hand stimuli) and then tested them on the other (e.g., predicted 1/3 of the Face stimuli). We first performed this analysis using a single PLSR component, and found a trend (Hand: *r*_*P(8)*_=0.281, *p*_*1*_=0.093; Face: *r*_*P(8)*_=0.3, *p*_*1*_=0.071). Using two components in the PLSR analyses improved results, which now became significant for the Hand (*r*_*P(8)*_=0.575, *p*_*1*_=9×10^−4^) and near significant for the Face (*r*_*P(8)*_=0.331, *p*_*1*_=0.058). Increasing to 3 or 4 components did not further improve this level of decoding. We thus report the results using 2 components (Fig. 4e), and used two components also for the cross-stimulus decoding, which turned out significant in both directions.

##### 4.1.5.10. Resting State Connectivity Analysis

To interrogate what connectivity profile is characteristic for electrode-pairs with high intensity coding, we used Neurosynth (Neurosynth.org) to extract a whole brain resting state connectivity map for the MNI location of each of the 85 contact-pairs in the insula (Supplementary File 1). Using SPM12 (https://www.fil.ion.ucl.ac.uk/spm/software/spm12/), we performed a regression analysis (general linear model) which included the 85 voxelwise resting state connectivity maps and two predictors: the correlation between power and rating for the Hand in the early window, and for the Face in the late window. Results were thresholded at p<0.001, corrected for multiple comparisons using family wise error correction at the cluster level. Results were then illustrated on an inflated cortical template provide in SPM12, and significant voxels were attributed to specific brain regions using the anatomy toolbox 3.0 (https://www.fz-juelich.de/inm/inm-7/EN/Resources/_doc/SPM%20Anatomy%20Toolbox_node.html).

##### 4.1.5.11. Spike Sorting and Selection of Responsive Single-units

Three patients had microwires (Behnke-Fried electrodes, Ad-Tech Medical; Fried et al., 1999) in the insula protruding from the electrode tip (plusses in Fig. 1c). Spikes were detected and sorted using Wave_Clus2 (Quiroga et al., 2004) In short, raw data was filtered between 300-3000 Hz. As per default settings, spike waveforms were extracted from 0.625 ms before to 1.375 ms after the signal exceeded a 5*noise threshold, where noise was the unbiased estimate of the median absolute deviation. Wave_Clus2 sorted and clustered the waveforms automatically and were manually checked by author RB. Clusters were excluded in which >2% of spikes were observed with an inter-spike interval <2 ms or with firing rate <1 Hz. To identify cells that responded to our stimuli, we used a Wilcoxon signed rank test comparing spike counts during baseline (−1 s to 0 s) against that during the pain period (1 s to 2 s) for Hand and Face trials together. Only cells that showed a response to the stimuli (*p*_*1*_<0.05), irrespective of pain intensity, were considered for further analysis.

Similar to LFP analyses, a cell was said to show intensity coding, if spike counts rank correlated positively with reported intensity. Because JASP includes Bayesian statistics using Kendall’s Tau but not Spearman r, we used the former to quantify evidence for or against intensity coding.

##### 4.1.5.12. Broadband Power Analysis in Microelectrodes

To explore whether intensity coding in cells and the broadband power (BBP, 20-190 Hz; Fig. 2a) from the same microwire were related, for the 10 microwires that yielded responsive neurons (whether these neurons showed intensity coding or not) we quantified the association between BBP averaged over the pain period (1-2 s) and intensity ratings (1-2, 3-4, 5-6, 7-8) using rank correlation coefficients separately for face and hand videos (again using Kendall’s Tau to provide BF_+0_ estimates). All 8 microwires protruding from the same electrode were first re-referenced to the microwire with the least spiking and lowest artefacts, yielding seven microwire recordings for each of the 4 electrode-tips with wires in the insula. Data were filtered to remove 50 Hz noise and harmonics at 100 and 150 Hz. Subsequently, they were separated into trials of 4 s (−1 s to 3 s relative to video onset), downsampled to 400 Hz and visually checked for artifacts. The time-frequency decomposition of power followed the same procedure as for the macro contact recordings. Finally, intensity coding at the level of spikes (i.e., *r*_*K*_(spikes,rating)) and BBP (*r*_*K*_(BBP,rating)) from the same wire were compared using a Kendall’s Tau coefficient.

### 4.2. fMRI Experiment

#### 4.2.1. Participants

Twenty-five healthy volunteers participated in the study. The full dataset of two participants was excluded from the analyses because head motions were above 3mm. Analyses were performed on the remaining twenty-three participants (13 females; mean age =28.76 years old +-SD=6.16). The study was approved by the local ethics committee of the University of Amsterdam (project number: 2017-EXT-8542).

#### 4.2.2. Stimuli and Procedure

The video pain rating task was performed as described above with the following differences. Each trial started with a grey fixation cross lasting 7-10 s, followed by a red fixation cross for 1 s, followed by the 2 s video, followed by a red fixation cross lasting 2-8 s, followed by the rating scale ranging from ‘not painful at all’ (‘0’) to ‘most intense imaginable pain’ (‘10’). The design also includes another condition we will not analyse here, in which participants viewed videos varying in color saturation and had to report on a scale from ‘not a change’ (‘0’) to ‘a very big change’ (‘10’). Participants were asked to provide a rating by moving the bar along the scale using two buttons for right and left (index and middle finger) and a third one for confirming their response (ring finger) using their left hand. The direction of the scale and the initial position of the bar was randomized in each trial. The videos used for the Face and Hand conditions for the electrophysiology and fMRI experiment were generated in the same way but were not identical.The task was split up into 46 blocks of 30 trials each, 2 blocks of electrical pain stimulations and, 2 blocks of mechanical slaps by a belt and 2 blocks of videos with changes in color saturation (presented in 46 separate fMRI acquisition runs). The last condition will not be discussed in this manuscript. Anatomical images were recorded between the fourth and fifth run of fMRI acquisition.

#### 4.2.3. Data Acquisition

MRI images were acquired with a 3-Tesla Philips Ingenia CX system using a 32-channel head coil. One T1-weighted structural image (matrix = 240×222; 170 slices; voxel size = 1×1×1mm) was collected per participant together with EPI (echo-planar imaging) volumes (matrix M x P: 80 × 78; 32 transversal slices acquired in ascending order; TR = 1.7 s; TE = 27.6 ms; flip angle: 72.90°; voxel size = 3×3×3 mm, including a .349 mm slice gap).

#### 4.2.4. Data Analysis

##### 4.2.4.1. Preprocessing

MRI data were processed in SPM12. EPI images were slice-time corrected to the middle slice and realigned to the mean EPI. High quality T1 images were coregistered to the mean EPI image and segmented. The normalization parameters computed during the segmentation were used to normalize the gray matter segment (1×1×1 mm) and the EPI images (2×2×2 mm) to the MNI templates. Finally, EPIs images were smoothed with a 6 mm kernel.

##### 4.2.4.2. Univariate regression

At the first level, we defined separate regressors for the Hand videos, Face videos, rating-scale and button-presses. Analyses focused on the 2 s that the videos were presented. The trial-by-trial ratings given by the participants were used as a parametric modulator, one modulator for the face and one for the Hand trials, on the respective video regressor. The rating-scale regressor started from the moment the scale appeared and ended with participants’ confirmation button press. The button-press regressor, finally, had zero duration and was aligned to the moment of all the button presses. Six additional regressors of no interest were included to model head movements. To quantify the degree to which each voxel in the insula had BOLD (blood-oxygen-level-dependent imaging) activity associated with trial-by-trial ratings, we then brought the parameter estimate for the parametric modulator obtained for Hand and Face trials separately to a second level t-test with 23 participants, and then used the resulting t-value as a measure of the random effect effect size of the association. We used t-values rather than the average value of the parametric modulator, because these values are to be compared against out-of-sample values of patients, and the topography of t-values is a better predictor for out-of-sample generalizations. However, the average parameter value correlated above 0.9 with the t-value.

##### 4.2.4.3. Multivariate regression

To investigate whether the pattern of BOLD activity across all voxels in the insula encodes intensity, we additionally performed a multivariate regression analysis akin to the PLSR for the BBP described in above. For each participant, we performed a general linear model that estimated a separate parameter estimate for the video epoch of trials in which participants gave a rating of 0-2, 3-4, 5-6 or 7-8 respectively, separately for Hand and Face trials. In matlab we then loaded for each subject the parameter estimate images for each level of rating, and only included voxels that fell within our insula mask (Faillenot et al., 2017). We then trained a weighted partial least-square regression using the matlab function plsregress and the data from all but one participant to predict rating based on a linear combination of the parameter estimates in each voxel and then used this linear combination to predict the rating of the left-out participant, then repeated the procedure for each participant. We weighted the regression by replicating each parameter estimate image in the training and testing set by the number of trials that went into it. Then we quantified how accurately the regression predicted the rating of the left-out participants using Kendall’s tau. We then tested whether the performance was above chance by comparing the 23 prediction accuracies (Kendall’s Tau) against zero in a one-tailed test. Based on the analysis on the BBP we performed this analysis with procedure using 2 or 3 components.

## Abbreviations

BA44/45: Brodmann Area 44/45
BBP: Broad-Band Power 20-190 Hz
BF_10_: Bayes Factor in favour of a two-tailed alternative hypothesis
BF_01_: Bayes Factor in favour of the null hypothesis relative to the two-tailed alternative hypothesis
BF_+0_: Bayes Factor in favour of a one-tailed alternative hypothesis
Fp1: Frontal pole region 1
iEEG: intracranial EEG
MNI: Montreal Neurological Institute coordinate system
OP8/9: Frontal Opercular regions 8 and 9
*p*_*1*_: one-tailed *p*-value
*p*_*2*_: two-tailed *p*-value
*r*_*P*_: Pearson’s correlation coefficient
*r*_*S*_: Spearman’s rank correlation coefficient
*r*_*K*_: Kendall’s Tau correlation coefficient.

## Data Availability

The data presented in this work is available at osf.io under the same title.

## Acknowledgments

We thank Pieter Roelfsema for enabling the collaboration that led to the access to the patients, Eline Ramaaker for her assistance in electrode localization, Agneta Fischer and George Bulte at UvA for advice and help with the use of FaceReader, and Tess den Uyl for advice on how to use FaceReader specifically to analyze facial expressions of pain.

