## Supplementary File 1 for "Intracranial Human Recordings Reveal Intensity Coding for the Pain of Others in the Insula"

| P ID | X | Y | Z |
|---|---|---|---|
| A | 28.5 | 12 | -12 |
| A | 29 | 15 | -8 |
| A | 29.5 | 17.5 | -4 |
| A | 30 | 20.5 | 0 |
| A | 30 | 23 | 4 |
| B | 38.5 | 6 | -12 |
| B | 38.5 | 8.5 | -7.5 |
| B | 37 | 11 | -2.5 |
| B | 36 | 14 | 2 |
| B | 36 | 17 | 6 |
| B | -41 | -5.5 | -7.5 |
| B | -40.5 | -2.5 | -2.5 |
| B | -39 | 0.5 | 2 |
| B | -37 | 3 | 7 |
| B | 40 | -4.5 | -12 |
| B | 39 | -7.5 | -7.5 |
| B | 37 | -10.5 | -2.5 |
| B | 36 | -13 | 2 |
| B | 36 | -15.5 | 6 |
| B | 35 | -18.5 | 10.5 |
| B | 33.5 | -21 | 15.5 |

| P ID | X | Y | Z |
|---|---|---|---|
| C | 38 | -1.5 | -8 |
| C | 38 | -4.5 | -4 |
| C | 38 | -7.5 | 0 |
| C | 38 | -10.5 | 5 |
| C | 37.5 | -13.5 | 10 |
| C | 36.5 | -16 | 14.5 |
| C | -34 | 2 | -19 |
| C | 39 | 2 | -15 |
| D | -35 | -6 | -6.5 |
| D | -35 | -7 | -1.5 |
| D | -35 | -8 | 3.5 |
| D | -34.5 | -9 | 9 |
| D | -33.5 | -10 | 14 |
| D | -37 | 12 | -8.5 |
| D | -36.5 | 15 | -4.5 |
| D | -36 | 18 | -1 |
| D | 40 | -5 | -13.5 |
| D | 39.5 | -7 | -8.5 |
| D | 39 | -9.5 | -4 |
| D | 38.5 | -12.5 | 0 |
| D | 38 | -15 | 4.5 |
| D | 38 | -17 | 9.5 |
| D | 38 | -19 | 14 |
| D | -37 | 8 | -14 |

| P ID | X | Y | Z |
|---|---|---|---|
| E | 38 | -5.5 | -6 |
| E | 38 | -8.5 | -2 |
| E | 37 | -11 | 2.5 |
| E | 36 | -14 | 7 |
| E | 35 | -17 | 11 |
| E | 34 | -20 | 15.5 |
| E | 40 | -1 | -14 |
| F | 38.5 | 5 | -13 |
| F | 39 | 7 | -8 |
| F | 39.5 | 9 | -3 |
| F | 40 | 11 | 2 |
| F | 36 | -9 | -5 |
| F | 36 | -11 | 0 |
| F | 36 | -13 | 5 |
| F | 36 | -14.5 | 10.5 |
| F | 36 | -16 | 15.5 |
| F | -33.5 | 12 | -13 |
| F | -32.5 | 16 | -8 |
| F | -32 | 21 | -4 |
| F | -32 | 26 | 0.5 |
| F | -36 | 3.5 | -15.5 |
| F | -35.5 | 2 | -11 |
| F | -35 | 0 | -4 |
| F | -35 | -2 | 3 |
| F | -35 | -4 | 9.5 |

| P ID | X | Y | Z |
|---|---|---|---|
| G | 40 | 6 | -8 |
| G | 39 | 7 | -2 |
| G | 38 | 9 | 3.5 |
| G | 39.5 | -3 | -8.5 |
| G | 38.5 | -6 | -4 |
| G | 37 | -9 | 0 |
| G | 35.5 | -11 | 4.5 |
| G | 34.5 | -13 | 9.5 |
| G | 34 | -15.5 | 15 |
| G | -38 | 1 | -10.5 |
| G | -37 | 3.5 | -5.5 |
| G | -35.5 | 6.5 | -1 |
| G | -34.5 | 9 | 3.5 |
| G | -33.5 | 11 | 9 |
| G | -36.5 | -11 | -7.5 |
| G | -35.5 | -13 | -2 |
| G | -35 | -14.5 | 3.5 |
| G | -35 | -16 | 9 |
| G | -34.5 | -18 | 15 |

**Supplementary File 1. MNI coordinates of recording sites.** In black, the MNI coordinates resulting from the average MNI coordinates of the two adjacent electrodes used for re-referencing for all macro electrodes. In gray the MNI coordinates of the micro electrodes. The color coding and patient identifiers reflect those used in Fig. 1c.
