## Supplementary File 2 for "Intracranial Human Recordings Reveal Intensity Coding for the Pain of Others in the Insula"

| Positive regression with intensity coding for Hand t=3.19, p<0.001, FWEc=772 | | | | | | | | | |
|---|---|---|---|---|---|---|---|---|---|
| Cluster size | Nr. vxls | % vxls assigned | Hem | Cito or anatamical region | % area activated | T-val | x | y | z |
| 9115 | 299.5 | 3.3 | L | Area Fp1 | 16.6 | 4.75 | -22 | 56 | -12 |
| | 206.8 | 2.3 | L | Area 44 | 23.6 | | | | |
| | 95.1 | 1 | L | Thal: Prefrontal | 15.1 | | | | |
| | 90.3 | 1 | L | Thal: Temporal | 17 | | | | |
| | 37.3 | 0.4 | L | Area 45 | 5.3 | 5.54 | -50 | 20 | -2 |
| | 23.6 | 0.3 | L | Area Fo2 | 2.3 | | | | |
| | 18.5 | 0.2 | R | Area 33 | 8.5 | | | | |
| | 14.3 | 0.2 | L | Area 33 | 6.7 | | | | |
| | 12.9 | 0.1 | R | Thal: Prefrontal | 2.3 | | | | |
| | | | L | IFG (p. Orbitalis) | | 5.78 | -34 | 30 | -8 |
| | | | | | | 5.78 | -38 | 30 | -6 |
| | | | | | | 5.73 | -44 | 22 | -6 |
| | | | R | Caudate Nucleus | | 4.81 | 20 | 24 | -2 |
| | | | L | Superior Orbital Gyrus | | 4.76 | -20 | 52 | -12 |
| 1590 | 432.1 | 27.2 | R | Lobule VIIa crusI (Hem) | 13.3 | 4.05 | 42 | -56 | -32 |
| | | | | | | 4.02 | 46 | -58 | -32 |
| | | | | | | 3.81 | 30 | -70 | -26 |
| | 394.8 | 24.8 | R | Lobule VI (Hem) | 21.9 | 3.98 | 32 | -58 | -28 |
| | | | | | | 3.77 | 24 | -66 | -30 |
| | 128.8 | 8.1 | R | Lobule VIIb (Hem) | 19.7 | 4.79 | 46 | -54 | -60 |
| | 73.8 | 4.6 | R | Lobule VIIa crusII (Hem) | 5.2 | | | | |
| | 64.1 | 4 | R | Area FG2 | 19.7 | | | | |
| | 32 | 2 | R | Lobule VI (Verm) | 13.8 | | | | |
| | 21.1 | 1.3 | R | Area hOc4v [V4(v)] | 3.4 | | | | |
| | 16 | 1 | R | Area FG1 | 6.4 | | | | |
| 772 | 249.6 | 32.3 | R | Area 45 | 24.2 | 4.68 | 60 | 24 | 0 |
| | | | | | | 3.67 | 64 | 20 | 18 |
| | | | R | IFG (p. Orbitalis) | | 4.46 | 42 | 26 | -10 |
| | | | | | | 4.44 | 44 | 28 | -8 |
| | | | | | | 4.02 | 58 | 38 | -8 |
| Negative regression with intensity coding for Hand t=3.19, p<0.001, FWEc=602 | | | | | | | | | |
| 2883 | 383.6 | 13.3 | R | Area hOc4la | 43.3 | 4.06 | 48 | -72 | -6 |
| | 208.3 | 7.2 | R | Area FG4 | 42.5 | 4.22 | 36 | -42 | -20 |
| | 207.6 | 7.2 | R | AreaFG3 | 31.7 | 4.21 | 40 | -40 | -24 |
| | 137.9 | 4.8 | R | Area hOc4v [V4(v)] | 22.2 | 4.55 | 46 | -82 | -20 |
| | 102.4 | 3.6 | R | Area hOc3v [V3v] | 12 | | | | |
| | 84.8 | 2.9 | R | Subiculum | 22.3 | 5.08 | 22 | -36 | -10 |
| | 82.1 | 2.8 | R | CA1 (Hippocampus) | 28.4 | | | | |
| | 81.5 | 2.8 | R | Amygdala (LB) | 38.2 | | | | |
| | 72.1 | 2.5 | R | Area FG2 | 22.2 | | | | |
| | 50.5 | 1.8 | R | Area hOc4lp | 9 | | | | |
| | 43.4 | 1.5 | R | Amygdala (SF) | 91.1 | | | | |
| | 40.6 | 1.4 | R | Lobule I IV (Hem) | 8.1 | 4.92 | 4 | -42 | -4 |
| | 27.6 | 1 | R | Thal: Parietal | 8.3 | | | | |
| | 24.5 | 0.8 | R | Area hOc5 [V5/MT] | 42.1 | | | | |
| | 19.8 | 0.7 | L | Lobule I IV (Hem) | 4.1 | 4.34 | -2 | -46 | -2 |
| | 18.5 | 0.6 | R | Thal: Temporal | 3.4 | | | | |
| | 14 | 0.5 | R | Lobule V (Hem) | 1.8 | | | | |
| | 12.9 | 0.4 | R | Area hOc1 [V1] | 0.6 | | | | |
| | 10.5 | 0.4 | R | HATA Region | 48 | | | | |
| | 10.4 | 0.4 | R | Area hOc2 [V2] | 1 | | | | |
| | 10.4 | 0.4 | R | Amygdala (CM) | 37.9 | | | | |
| | | | R | Lingual Gyrus | | 4.2 | 22 | -86 | -4 |
| 2277 | 363.9 | 16 | R | Area 4a | 33.4 | 3.41 | 30 | -34 | 70 |
| | 137.5 | 6 | R | Area 3b | 21.9 | 3.39 | 34 | -34 | 58 |
| | 114.9 | 5 | L | Area 1 | 20.2 | | | | |
| | 108.9 | 4.8 | R | Area 1 | 15.5 | | | | |
| | 74.8 | 3.3 | L | Area 5L (SPL) | 10.8 | | | | |
| | 71.6 | 3.1 | L | Area 4a | 7.7 | | | | |
| | 66.1 | 2.9 | L | Area 5M (SPL) | 13.6 | | | | |
| | 60.9 | 2.7 | R | Area 5L (SPL) | 8.3 | | | | |
| | 38.5 | 1.7 | L | Area 2 | 7.3 | | | | |
| | 17.6 | 0.8 | R | Area 5M (SPL) | 6 | | | | |
| | 17.4 | 0.8 | R | Area 2 | 2.7 | | | | |
| | 16.4 | 0.7 | L | Area 3b | 2.9 | | | | |
| | 14.9 | 0.7 | R | Area 4p | 4.8 | | | | |
| | | | R | Paracentral Lobule | | 4.48 | 4 | -38 | 80 |
| | | | | | | 3.61 | 2 | -18 | 80 |
| | | | R | Posterior-Medial Frontal | | 3.42 | 14 | -10 | 68 |
| | | | L | Postcentral Gyrus | | 3.41 | -22 | -36 | 82 |
| 626 | 14.3 | 2.3 | R | Area OP3 [VS] | 6.8 | | | | |
| | 7.5 | 1.2 | R | Area 3a | 3.7 | | | | |
| | 3.8 | 0.6 | R | Area 3b | 0.6 | | | | |
| | 0.1 | 0 | R | Area OP1 [SII] | 0 | | | | |
| 604 | 319.1 | 52.8 | L | Area hOc4la | 37.3 | 3.77 | -42 | -82 | -16 |
| | | | | | | 3.73 | -48 | -78 | 0 |
| | 42.4 | 7 | L | Area FG2 | 8.3 | 3.66 | -42 | -74 | -12 |
| | | | | | | 3.64 | -38 | -72 | -8 |
| | 28.9 | 4.8 | L | Area hOc4v [V4(v)] | 4 | | | | |
| | 23.8 | 3.9 | L | Area hOc5 [V5/MT] | 29.5 | 3.66 | -40 | -74 | -4 |
| | 10.4 | 1.7 | L | Area hOc4lp | 1.2 | | | | |

**Supplementary File 2. Resting state connectivity results.** The table indicates for each significant cluster, the size in number of voxels of the cluster; the number of voxels of that clusters that have been assigned to cytoarchitectonic areas based on the Anatomy toolbox for SPM (http:// www.fz-juelich.de/ime/spm_anatomy_toolbox); the number of voxels assigned expressed in percentage; the hemisphere covered by the cluster (L=left, R=right); the cito-architectonic area those voxels have been assigned to when the information is available, or the anatomical description of the area voxels fell in; the percentage of the cytoarchitectonic area covered by the assigned voxels; the t-values of the identified peaks of activity with the respective MNI coordinates.
